# Single cell RNA sequencing redefines the mesenchymal cell landscape of mouse endometrium

**DOI:** 10.1101/2020.08.20.257246

**Authors:** PM Kirkwood, DA Gibson, JR Smith, JR Wilson-Kanamori, O Kelepouri, A Esnal-Zufiaurre, R Dobie, NC Henderson, PTK Saunders

## Abstract

The endometrium is a dynamic tissue that exhibits remarkable resilience to repeated episodes of differentiation, breakdown, regeneration and remodelling. Endometrial physiology relies on a complex interplay between the stromal and epithelial compartments with the former containing a mixture of fibroblasts, vascular and immune cells. There is evidence for rare populations of putative mesenchymal progenitor cells located in the perivascular niche of human endometrium, but the existence of an equivalent cell population in mouse is unclear.

In the current study we used the *Pdgfrb*-BAC-eGFP transgenic reporter mouse in combination with bulk and single cell RNA sequencing (scRNAseq) to redefine the endometrial mesenchyme. Contrary to previous reports we show that CD146 is expressed in both PDGFRβ+ perivascular cells as well as CD31+ endothelial cells. Bulk RNAseq revealed cells in the perivascular niche which express high levels of *Pdgfrb* as well as genes previously identified in pericytes and/or vascular smooth muscle cells (*Acta2, Myh11, Olfr78, Cspg4, Rgs4, Rgs5, Kcnj8, Abcc9*). scRNAseq identified five subpopulations of cells including closely related pericytes/vascular smooth muscle cells and three subpopulations of fibroblasts. All three fibroblast populations were PDGFRα+/CD34+ but were distinct in their expression of *Spon2/Angptl7* (fibroblast 1), *Smoc2/Rgs2* (fibroblast 2) and *Clec3b/Col14a1/Mmp3* (fibroblast 3), with potential functions in regulation of immune responses, response to wounding and organisation of extracellular matrix respectively.

In conclusion, these data are the first to provide a single cell atlas of the mesenchymal cell landscape in mouse endometrium. By identifying novel markers for subpopulations of mesenchymal cells we can use mouse models investigate their contribution to endometrial function, compare with other tissues and apply these findings to further our understanding of human endometrium.

**Highlights:**

- GFP expression in the mouse endometrium, under the control of the *Pdgfrb* promoter, is restricted to two cell populations based on the intensity of GFP with GFP^bright^ cells close to the vasculature
- Single cell RNAseq identified five subpopulations of GFP+ mesenchymal cells: pericytes, vascular smooth muscle cells (vSMC) and three closely related but distinct populations of fibroblasts
- Bioinformatics revealed that pericytes and vSMC share functions associated with the circulatory system, actin-filament process and cell adhesion, and an apparent role for pericytes in smooth muscle cell migration and response to interferons
- Comparisons between the fibroblast subpopulations suggest distinct roles in regulation of immune response, response to wound healing and collagen organisation.

**Graphical Abstract:** 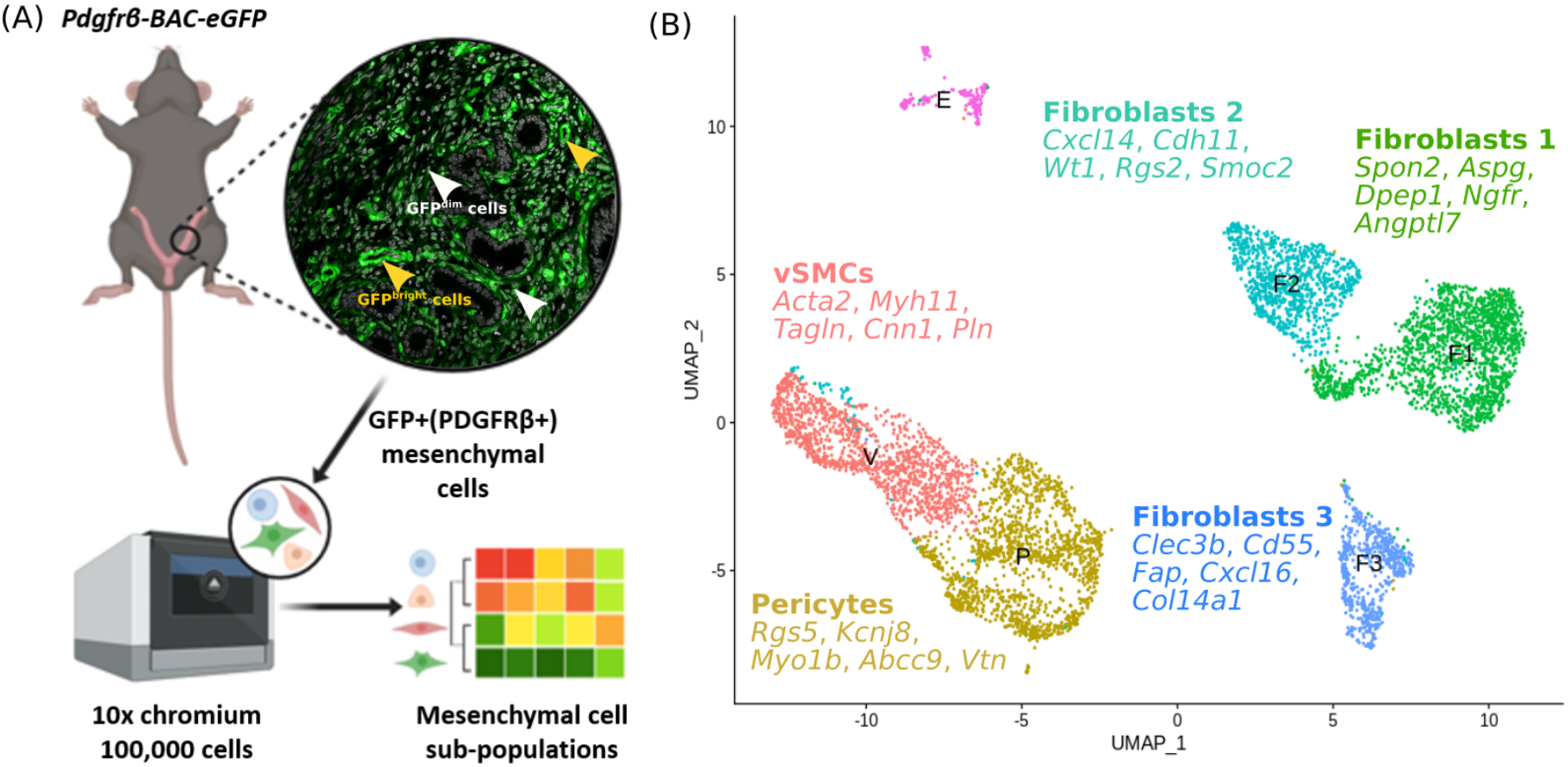

## INTRODUCTION

The endometrium of women and mice share a similar architecture with epithelial cells lining the glands and lumen supported on a complex stroma containing fibroblasts and a well-developed vasculature. In both species stromal fibroblasts transform into decidual secretory cells capable of supporting an implanting blastocyst in response to the action of the sex steroid hormones oestrogen and progesterone; such decidualization, either spontaneous (as in women) or requiring additional blastocyst-derived signals (as in mice), results in irreversible changes in stromal cell function (Gellersen, et al. 2007; Matsumoto 2017). In species with spontaneous decidualization in the absence of a viable blastocyst the region of tissue containing the decidual cells breaks down and is shed during menstruation (Bellofiore, et al. 2018b; Garry, et al. 2009; Maybin and Critchley 2015). Menstruation only occurs in women, and a few other species including higher primates and the Spiny mouse (*Acomys cahirinus*; (Bellofiore, et al. 2018a) and is characterised by spontaneous transformation/decidualization of stromal fibroblasts. A menstrual-like event can be simulated in other mouse species following artificial induction of decidualization (Cousins, et al. 2016; Cousins, et al. 2014).

In both natural and induced menstrual cycles the surface of the endometrium resembles a bloody wound during tissue breakdown but repair is both rapid and scar-free with evidence of epithelial cell proliferation, mobilisation of stromal cells and evidence of mesenchymal to epithelial cell transformation (Cousins et al. 2014; Garry et al. 2009). It has been postulated that the rapid restoration of endometrial tissue integrity is in part due to mobilisation and differentiation of tissue-resident endometrial progenitor cells (Gargett, et al. 2016). Many studies have searched for candidate endometrial progenitor cells using *in vitro* assays based on clonogenicity and self-renewal as well as whole tissue analysis including recovery of side population cells (SP) and those labelled with BrdU in label retention assays (Gargett and Masuda 2010; Tempest, et al. 2018). In 2004, Chan *et al*, reported the existence of small populations of epithelial and stromal cells in human endometrium that were able to form colonies from single cell suspensions *in vitro* (0.22% and 1.25% respectively, (Chan, et al. 2004). These cells also exhibited key features of stem-like cells including self-renewal, a high proliferative potential and the capacity for multilineage differentiation (Gargett, et al. 2009). A population of perivascular cells from human endometrium with clonogenic and multi-lineage differentiation potential *in vitro* was initially isolated based on co-expression of PDGFRβ and CD146 (Chan et al. 2004; Schwab, et al. 2005) but in further studies Sushi Domain Containing protein 2 (SUSD2), recognised by the W5C5 monoclonal antibody has emerged as a definitive candidate marker for putative endometrial mesenchymal progenitors (Gargett et al. 2016). This antibody reacts with human and primate proteins but not ovine (Letouzey, et al. 2015) or mouse. Human SUSD2+ cells exhibited clonogenicity, multilineage differentiation and self-renewal, representing ~4% of endometrial stromal cells with a perivascular location (Gargett et al. 2016; Masuda, et al. 2012).

Studies in mice have tried to identify tissue resident endometrial progenitor cells. For example, labelling of endometrial tissue with BrdU injections in 3-day old mice resulted in detection of LRCs in the stromal compartment into adulthood, falling from 8% of all stromal cells on day 49 to ~2% on day 112 post injection. Some of the LRCs expressed putative ‘stem cell’ markers Oct4 and c-kit/CD117 (Cervello, et al. 2007) as well as CD44, CD90, PDGFRβ, CD146 and Sall4 (Chan and Gargett 2006). Stromal LRC have a close association with CD31+ endothelial cells and expression of α-SMA appears consistent with a vascular smooth muscle cell and/or pericyte phenotype (Chan and Gargett 2006; Kaitu’u-Lino, et al. 2012). Although putative endometrial mesenchymal progenitor cells in the mouse are reported to co-express PDGFRβ and CD146 (Chan and Gargett 2006; Parasar, et al. 2017) to date no single marker equivalent to human SUSD2 has been described.

In summary, studies on human endometrial tissue suggest the stroma contains one or more populations of tissue resident progenitor cells that contribute to resilience of the tissue. Whilst candidate cells adjacent to the vasculature have been identified in human endometrium the evidence for their existence in mouse endometrium is less robust. The current study used the *Pdgfrb*-BAC-eGFP transgenic reporter mouse in combination with single cell transcriptomics to generate a definitive set of cell-specific markers for mesenchyme-derived cell populations. Not only did these studies identify and characterise a population of perivascular cells in the mouse endometrium but novel findings revealed evidence of unexpected complexity in the fibroblast populations that make up the bulk of the endometrial stromal mesenchyme.

## RESULTS

### Two distinct populations of PDGFRβ+ mesenchymal cells can be detected in the *Pdgfrb*-BAC-eGFP mouse endometrium based on relative expression of GFP and perivascular marker CD146

In the uterus of *Pdgfrb*-BAC-eGFP mice expression of GFP protein was detected in mesenchymal cells but not in epithelial cells (glandular: GE; luminal: LE) or smooth muscle cells of the myometrium (M) (Figure 1A). In tissue sections the intensity of GFP expression revealed two subpopulations of cells which we classified as GFP^dim^ and GFP^bright^ (Figure 1A; white and yellow arrow heads respectively). Co-localisation of GFP with PDGFRβ was confirmed in both GFP^dim^ and GFP^bright^ cells (Figure 1B; white and yellow arrow heads respectively). Sections were co-labelled with markers of epithelial (EpCAM), mesenchymal (desmin) and vascular (CD31: endothelial; CD146: endothelial/pericyte) cells. Both GFP+ populations were EpCAM- (Figure S1B(ii)) and desmin+ (Figure S1B(iii)). Notably GFP^dim^ cells were CD31-CD146- and GFP^bright^ cells were CD31-CD146+: the latter were located in close proximity to CD31+CD146+ endothelial cells (Figure 1C, split channel images Figure S1C (i-iiii)).

**Figure 1.**
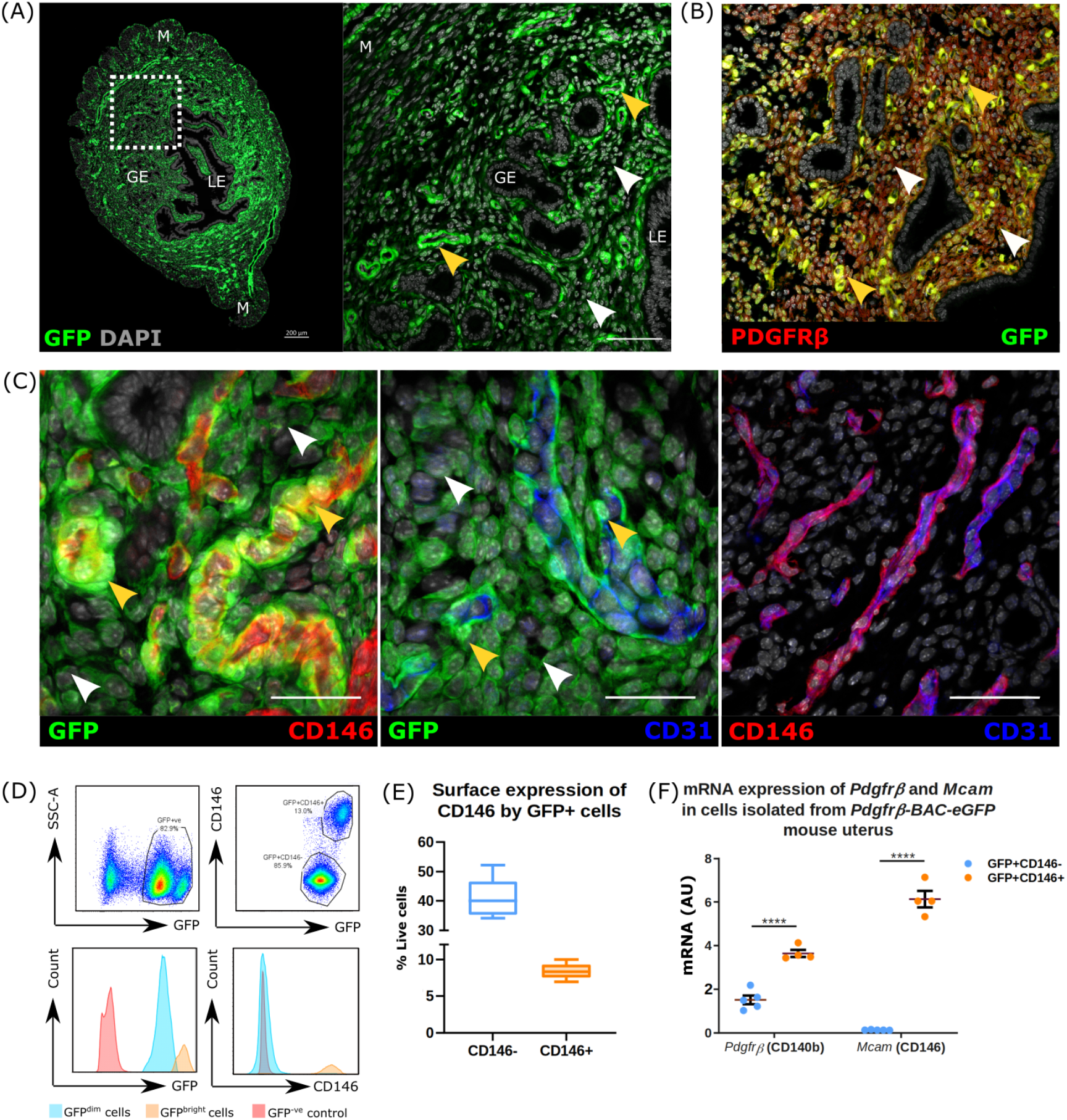
Expression of reporter protein (GFP) in the Pdgfrb-BAC-eGFP mouse endometrium identifies two distinct populations of mesenchymal cells. **(A)** Localisation of GFP+ cells in Pdgfrb-BAC-eGFP mouse endometrium by immunofluorescence identifies two populations of PDGFRβ expressing cells based on the intensity of GFP staining: GFP^dim^ cells (white arrow heads) and GFP^bright^ cells (yellow arrow heads). GFP reporting in the endometrium of the Pdgfrb-BAC-eGFP mouse is strictly mesenchymal, absent from both luminal (LE) and glandular epithelial cells (GE) and smooth muscle cells of the myometrium (M). **(B)** Immunofluorescence detection was used to confirm that expression of GFP efficiently reports that of native PDGFRβ protein whereby dual expression is detected in mesenchymal cells only. **(C)** Further immunofluorescence detection of GFP, endothelial cell marker CD31, and putative pericyte marker CD146 revealed that GFP^bright^ cells co-expressed CD146 and are localised adjacent to CD31+ endothelial cells in the perivascular niche, whereas GFP^dim^ expressing cells showed a wider distribution throughout the endometrial stroma and did not co-express CD146. Importantly endothelial cells were found to express both CD31 and CD146. **(D)** Flow cytometry analysis of Pdgfrb-BAC-eGFP mouse endometrium phenotypes two populations of live+/CD31-/CD45-/GFP+ cells based on the intensity of fluorescence: GFP^dim^ and GFP^bright^ cells. GFP+ cells can be further stratified based on their relative expression of CD146: GFP^dim^CD146-cells and GFP^bright^CD146+ cells. **(E)** Quantification of FC analysis reveals relative proportions of GFP^dim^CD146-cells (40.05±2.25% live cells; 82.78±1.29% GFP+ cells) and GFP^bright^CD146+ cells (8.33±0.36% live cells; 17.22±1.26% GFP+ cells) in the mouse uterus; n=8. **(G)** qPCR analysis of Pdgfrb and Mcam (CD146) mRNA expression in GFP^dim^CD146- and GFP^bright^CD146+ cells isolated from Pdgfrb-BAC-eGFP mouse uterus. GFP^bright^CD146+ cells expressed significantly higher mRNA levels of Pdgfrb and Mcam (CD146) than GFP^dim^CD146-cells (FC to Actb); n=4 (paired t-test, ****p<0.0001).

Histological findings were confirmed by FACS analysis of *Pdgfrb*-BAC-eGFP uterine tissues. After negative gating for endothelial (CD31+) and immune (CD45+) cells (Figure S2A(i-vi), Figure S2B) the total GFP+ population separated into two subpopulations based on GFP fluorescence intensity (GFP^dim^ and GFP^bright^; Figure 1D). CD146 expression followed histological findings: 40.05±2.25% live cells were GFP^dim^CD146- and 8.33±0.36% live cells were GFP^bright^CD146+ (Figure 1D-E). Immunohistochemistry on cytospins of FACS-sorted GFP+ subpopulations confirmed that GFP^dim^ cells were GFP+PDGFRβ+CD146-while GFP^bright^ cells were GFP+PDGFRβ+CD146+ (Figure S1E), and that GFP^bright^ cells had significantly higher expression of *Pdgfrb* and *Mcam* mRNAs than GFP^dim^ cells (Figure 1F).

Collectively these results phenotype two subpopulations of GFP+/PDGFRβ expressing cells in the uterus of the *Pdgfrb*-BAC-eGFP mouse as GFP^dim^CD146-cells and GFP^bright^CD146+ cells.

### Bulk mRNA sequencing of uterine mesenchymal subpopulations from the *Pdgfrb*-BAC-eGFP mouse identifies a transcriptomic signature characteristic of perivascular cells

To investigate the transcriptomic profile and potential functional role of perivascular cells in the endometrium, GFP^bright^ and GFP^dim^ cells were isolated from *Pdgfrb*-BAC-eGFP uterine tissues and compared using bulk mRMA sequencing. The concentration of purified RNA from GFP^bright^ cells was insufficient for downstream sequencing applications due to a lower number of sorted cells. Therefore, GFP^total^ cells (GFP^dim^ + GFP^bright^) were compared against GFP^dim^ cells to infer the profile of GFP^bright^ cells (GFP^total^-GFP^dim^).

PCA analysis revealed that the two sample groups cluster separately, indicating distinct transcriptomic signatures that we attributed to the presence of GFP^bright^ cells in the GFP^total^ samples (Figure 2A). Differential gene expression analysis revealed 143 genes significantly upregulated in GFP^total^ samples when compared to GFP^dim^ samples (as determined by FDR). Amongst the most significantly upregulated were key genes *Acta2, Kcnj8, Olfr78, Rgs4, Rgs5, Myh11, Abcc9, Cspg4, Sox6, Myom1, Ednra, Ednrb, Kitl, Myo1b, Olfr78, Olfr558* and *Vcam1* (Figure 2B-C). Notably a clear increase in the mean CPM and LogFC of *Pdgfrβ* and *Mcam* (CD146) was confirmed in GFP^total^ samples when compared to GFP^dim^ samples (Figure 1D), consistent with the expression of such genes by GFP^bright^ cells. The resultant GFP^bright^ cell signature was enriched for gene ontology (GO) biological process (BP) terms relating to blood circulation, muscle cell differentiation, leukocyte migration, regulation of blood pressure and smooth muscle contraction among others (Figure 2E). Further analysis of specific GO terms associated *Cxcl10, Cxcl11, Cxcl12, Csf1, Itga2, Itga4, Ednra* and *Icam1* with the regulation of leukocyte migration and *Hrc, Ptgs2, Trpm4, Cnn1, Myocd, Tpm1, Pln* and *Acta2* with the regulation of smooth muscle cell contraction (Figure 2F, Figure S3A-B). Gene network analysis also highlighted core genes (leading edge subset) that were associated with multiple GOBP terms and might therefore have more prominent functional significance *in vivo*. These included *Kcnj8, Acta2, Ptgs2, Rgs4, Icam1, Cxcl10, Ednra* and *Ednrb* (Figure S3C).

**Figure 2.**
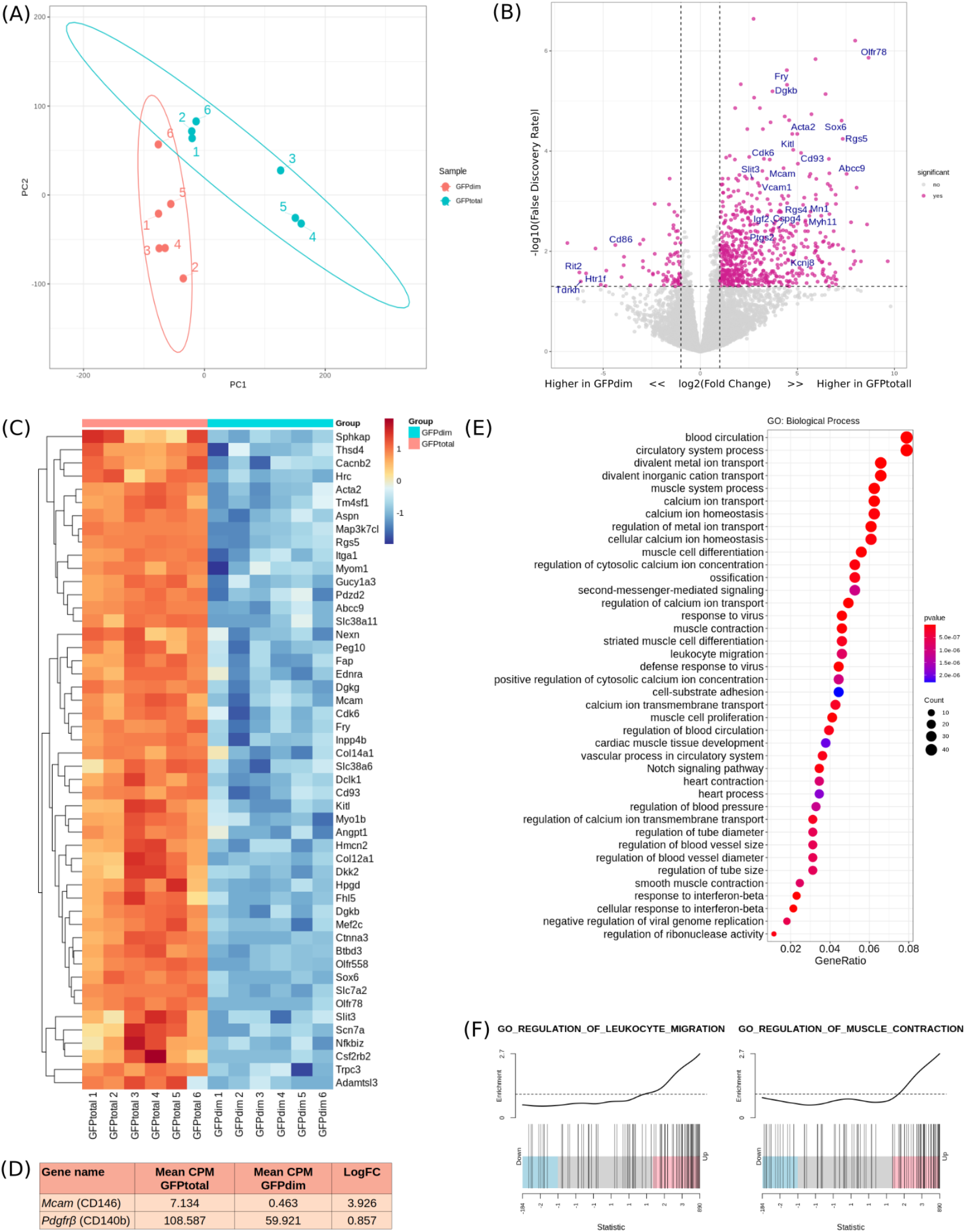
Bulk RNA sequencing of GFP^total^ and GFP^dim^ cells isolated from Pdgfrb-BAC-eGFP mouse endometrium infers a transcriptomic signature of GFP^bright^ cells characteristic to that of perivascular cells. **(A)** Principal component plot of the first and second components from PCA analysis using selected GFP^total^ and GFP^dim^ samples (n=6). **(B)** Volcano plot: differential gene expression between groups GFP^total^ and GFP^dim^ defining significantly up- or down-regulated genes as those with Log_2_FC>2 and log_10_FDR>1. **(C)** Heatmap: top 50 most differentially expressed genes between GFP^total^ and GFP^dim^ cells as determined by FDR. **(D)** Table outlining the mean CPM and LogFC of target genes Pdgfrb and Mcam (CD146). **(E)** Dotplot: gene ontology (GO) analysis of the putative perivascular cell genes to identify key biological processes (BP) attributed to the inferred GFP^bright^ cells transcriptome; dot size: number of genes in data attributed to each GO term; dot colour: pvalue representing enrichment score. **(F)** Enrichment plots: distribution of putative perivascular cell genes associated with ‘regulation of leukocyte migration’ and ‘regulation of smooth muscle cell contraction’; line peak represents the enrichment score; blue: genes under-represented in data; pink: genes over-represented in data.

Interestingly, genes associated with GFP^bright^ cells including those commonly expressed by vascular smooth muscle cells (vSMCs; *Acta2, Myh11, Pln*) as well as those commonly expressed by pericytes (*Kcnj8*, *Rgs5*, *Abcc9*). GO analysis also identified cellular function often associated with these two cell types including blood vessel regulation, cell adhesion, cell contraction and immune responses. These results suggest that co-expression of *Pdgfrb* and *Mcam* (CD146) identifies a heterogeneous population of perivascular cells (GFP^bright^ cells) which may include both vSMCs and pericytes.

### Validation of genes enriched in GFP^bright^ cells

Validation of 10 putative GFP^bright^-specific genes (LogFC>2, FDR<0.05, Figure S4A-B) was performed by qPCR analysis of mRNA expression in a matched set of samples (GFP^total^ and GFP^dim^ cells, Figure S4C) and an independent cohort of GFP^dim^CD146- (fibroblasts), GFP^bright^CD146+ (perivascular) and CD31+ (endothelial) cells isolated from *Pdgfrb*-BAC-eGFP uterine tissues (Figure 3A).

**Figure 3.**
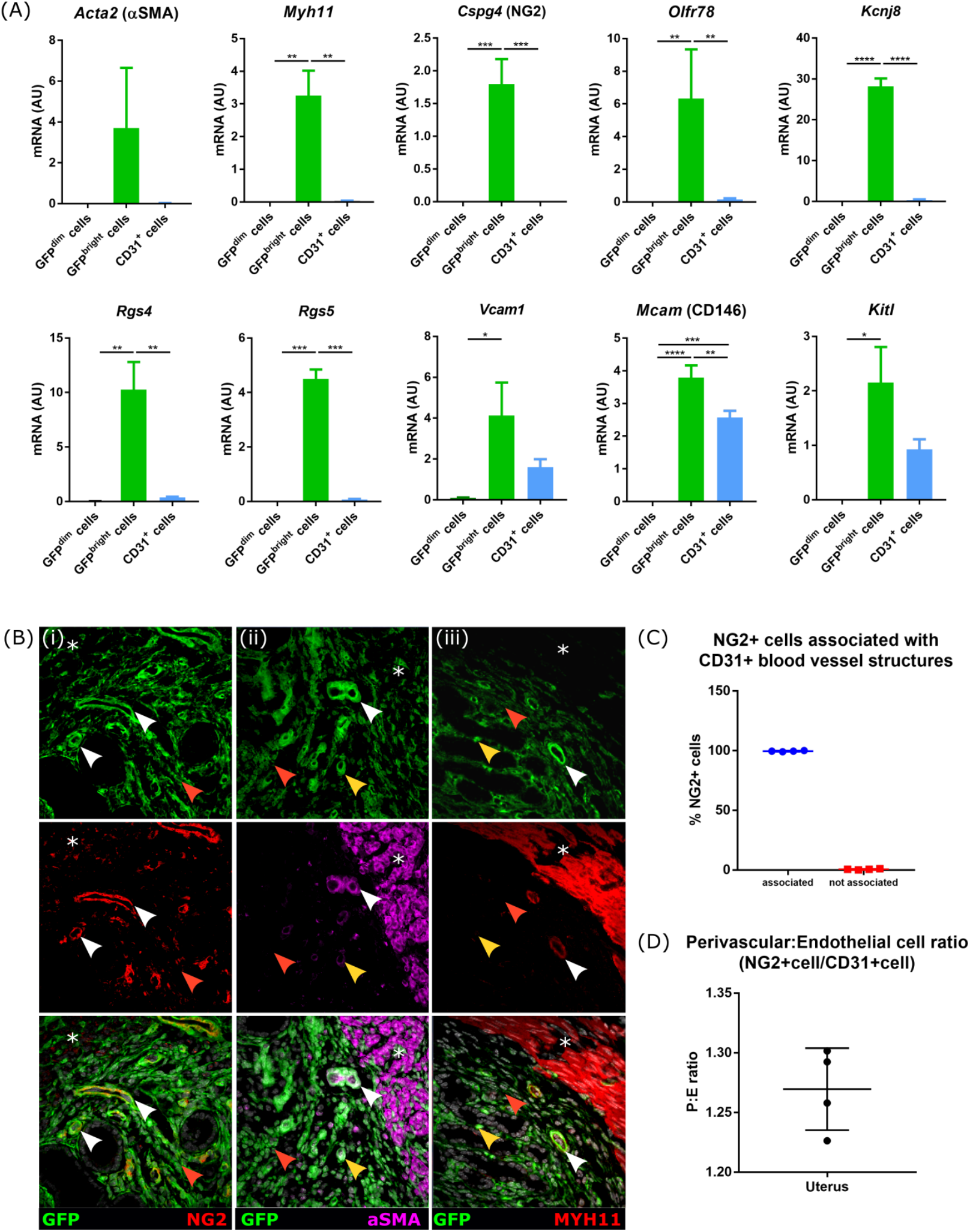
Validation of Bulk mRNA sequencing results in mouse uterine tissue samples using both qPCR and immunofluorescence. **(A)** qPCR gene expression analysis of perivascular cell genes in GFP^dim^ cells (n=3), GFP^bright^ cells (n=3) and CD31+ cells (n=4) isolated from Pdgfrb-BAC-eGFP uterine tissue. mRNA for Acta2, Myh11, Cspg4 (NG2), Olfr78, Rgs4, Rgs5, Vcam1, Mcam (CD146), Kitl and Kcnj8 was detected in GFP^bright^ cells and not in GFP^dim^ cells while mRNA for Vcam1, Mcam (CD146) and Kitl were also detected in CD31+ cells (oneway ANOVA, Holm-Sidak’s multiple comparisons test, *p<0.05, **p<0.01, ***p<0.001, ****p<0.0001). **(B)** Analysis of GFP, NG2 (Cspg4), αSMA and MHY11 in Pdgfrb-BAC-eGFP and C57BL/6 (wild type) murine uterine tissue by immunofluorescence: **(I)** NG2 expression was detected in all perivascular GFP^bright^ cells and completely absent from GFP^dim^ cells. **(ii)** αSMA expression was detected in a subset of GFP^bright^ cells most predominantly located at the endometrial/myometrial junction (white arrow heads). GFP^bright^ cells more proximal to the lumen did not coexpress αSMA (yellow arrow heads) nor did GFP^dim^ cells (red arrow heads). **(iii)** Similarly, MYH11 expression was only detected in subset of GFP^bright^ cells located at the endometrial/myometrial junction (white arrow heads) not those more proximal to the lumen (yellow arrow heads) or GFP^dim^ cells (red arrow heads). Smooth muscle cells of the myometrium expressed both αSMA and MYH11 but not NG2 ((i-iii) asterisk) (representative images; n=4). **(C-D)** Quantification of NG2+ perivascular and CD31+ endothelial cells mouse uterine tissue sections: (C) number of NG2+perivascular cells associated with CD31+ blood vessel structures; (D) ratio of number of NG2+ perivascular cells to CD31+ endothelial cells (n=4).

All putative perivascular cell-associated genes were substantially expressed in GFP^bright^CD146+ cells, and no expression was detected for any in GFP^dim^CD146-cells (Figure 3A). CD31+ cells expressed substantial levels of *Vcam1, Mcam*, and *Kitl*, and lower expression of *Olfr78*, *Rgs4*, *Rgs5*, and *Kcnj8*, but in all cases expression was significantly higher in GFP^bright^CD146+ cells (Figure 3A). This provided further validation of the strategy employed to identify a perivascular cell gene signature.

### Identification of proteins associated with the GFP^bright^ cell signature in tissue sections

Immunoexpression of *Cspg4* (NG2), *Acta2* (αSMA) and *Myh11*, revealed NG2 was expressed ubiquitously and exclusively by GFP^bright^ cells in endometrium (Figure 3B(i); white arrow heads). Interestingly, αSMA (Acta2) was detected in a subset of GFP^bright^ cells located predominantly at the endometrial/myometrial junction in the basal endometrium (Figure 3B(ii); white arrow heads) and the GFP^bright^ cells more proximal to the endometrial lumen did not express αSMA (Figure 3B(ii); yellow arrow heads). A similar pattern of expression was observed when examining MYH11 (Figure 3B(iii)). Importantly, NG2, αSMA and MYH11 were not detected in GFP^dim^ cells (Figure 3B(i-iii); red arrow heads) but αSMA and MYH11 were both present in smooth muscle cells of the myometrium (Figure 3B(ii-iii); asterisk). These results confirmed heterogeneity within the GFP^bright^ cell population dependent upon location: αSMA+/MYH11+/NG2+ cells located around basal blood vessels and were likely vSMCs, while αSMA-/MYH11-/NG2+ cells were more proximal to the lumen and likely represent pericytes. The functional significance of such defined topography and whether this relates to different types of blood vessels remains to be investigated.

Only NG2 expression was specific/exclusive to perivascular cells in the mouse endometrium and was therefore used to determine the relationship between perivascular cells (NG2+ cells with visible nucleus) and endothelial cells (CD31+ cells with visible nucleus). All NG2+ cells were located in close proximity to CD31+ endothelial cells (100±0.24%; Figure 3C) and the ratio NG2:CD31 expressing cells was calculated to be 1.25:1 (±0.02) (Figure 3D).

### Deconvolution of the mouse uterus by single-cell RNA sequencing identifies five distinct subpopulations of mesenchymal cells present in the endometrium during homeostasis

Bulk RNA sequencing identified a transcriptomic profile of GFP^bright^ cells similar to that of both vSMCs and pericytes suggesting heterogeneity within this cell population. To extend these findings and analyse the full range of cells within the uterine mesenchyme, we performed single-cell RNA sequencing (scRNAseq) on the GFP+ cell fraction of the *Pdgfrb*-BAC-eGFP mouse uterus (n=4). Sequencing detected 6,379 individual cells with 71,067 mean reads per cell equating to 1,847 median genes per cell.

Unsupervised clustering based on principal components of the most variably expressed genes partitioned cells into 6 individual clusters (Figure 4A). Expression of signature genes associated with known endometrial cell phenotypes identified five of the six clusters as mesenchymal cells (positive for *Pdgfrb, Vim* and *Des*; Figure 4C(i)), two of which showed preferential expression of canonical perivascular markers including *Mcam* (CD146), *Acta2* (αSMA) and *Cspg4* (NG2) (Figure 4C(ii)), and the remaining three showing preferential expression of canonical fibroblast markers such as *Cd34*, *Pdgfra* and *Mfap5* (Figure 4C(iii)). The final cluster was found to express genes typical of epithelial cells (*Cdh1, Epcam, Krt18*; Figure 4C(iv)). We labelled the six subpopulations as *Pdgfrb+Mcam*-fibroblasts (F1, F2, F3), *Pdgfrb+Mcam+* perivascular cells (P; pericytes, V; vSMCs) and a small population of *Pdgfrb-Mcam*-epithelial cells (E) suspected to represent contamination at the time of cell purification.

**Figure 4.**
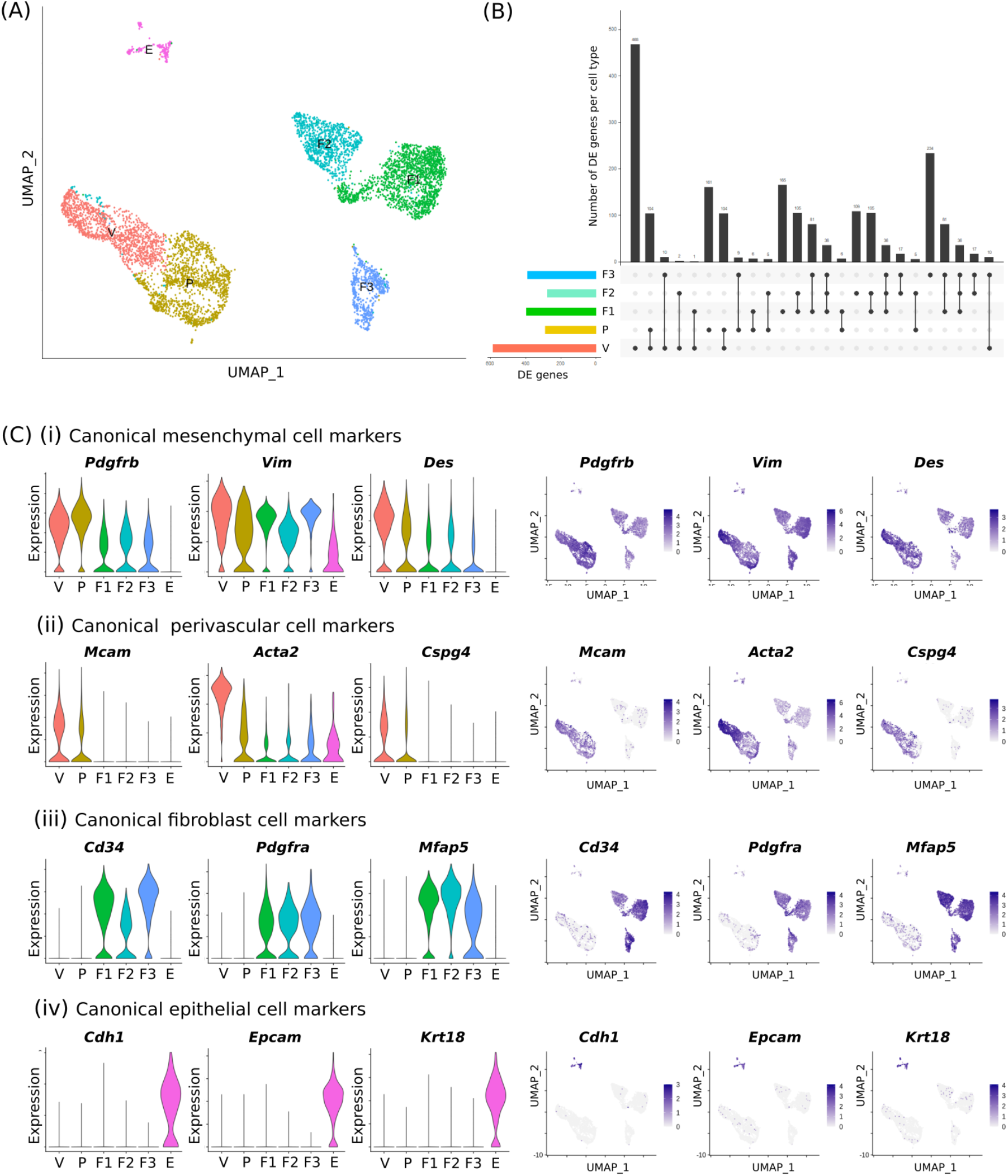
Single-cell RNA sequencing analysis of total GFP+ cells isolated from Pdgfrb-BAC-eGFP mouse uterine tissue reveals a previously unidentified heterogeneity within the murine endometrial mesenchyme during homeostasis. **(A)** Uniform manifold approximation and projection (UMAP) visualisation: 6,379 GFP+ mesenchymal cells isolated from Pdgfrb-BAC-eGFP mouse uterine tissue (n=4; median nGENE= 1847, nUMI= 4316) cluster into six distinct subpopulations. **(B)** Upset plot: visualisation of the intersection between gene lists for each cell cluster: each bar represents a set of genes; each row represents a cell cluster; individual dots represent a gene set unique to that cell cluster while connected dots represent gene sets that are shared between cell clusters. The top five intersections per cluster were evaluated. **(C)** (i-iv) Violin plots and UMAP visualisations: expression of gene signatures associated with known cell types present in the endometrium used to infer cell lineages for each cell cluster: (i) canonical mesenchymal cell markers Pdgfrb, Vim and Des; (ii) canonical perivascular cell markers Mcam (CD146), Acta2 (αSMA) and Cspg4 (NG2); (iii) canonical fibroblast cell markers Cd34, Pdgfra and Mfap5; and (iv) canonical epithelial cell markers Cdh1, Epcam and Krt18. Expression patterns identified three populations of stromal fibroblasts, two populations of perivascular cells (pericytes, vascular smooth muscle cells (vSMCs)), and a small contaminating population of epithelial cells. Fibroblasts 1 (F1), 2 (F2) & 3 (F3) to have low expression of Pdgfrb (representative of GFP^dim^ fraction) while pericytes & vSMCs have high expression of Pdgfrb and co-expression of Mcam (representative of GFP^bright^ fraction). Confirms capture of both subpopulations of mesenchymal cells described through previous methods.

Differential gene expression analysis revealed a high degree of similarity between putative vSMCs (V) and pericytes (P) and between the three fibroblast clusters (F1, F2, F3) (Figure 4B, Figure 5A). Importantly, a unique signature can be deciphered for each cell cluster revealing target genes specifically attributed to each individual cell type. Genes expressed preferentially by vSMCs include *Myh11, Tagln, Cnn1* and *Kitl* (Figure 5B(i)); pericytes include *Kcnj8, Myob1, Abcc9* and *Ednrb* (Figure 5B(ii)); F1 include *Aspg, Dpep1, Ngfr* and *Angptl7* (Figure 5B(iii)); F2 include *Cxcl14, Cdh11, Wt1* and *Rgs2* (Figure 5B(iv)); and F3 include *Clec3b, Fap, Cd55* and *Vit* (Figure 5B(v)).

**Figure 5.**
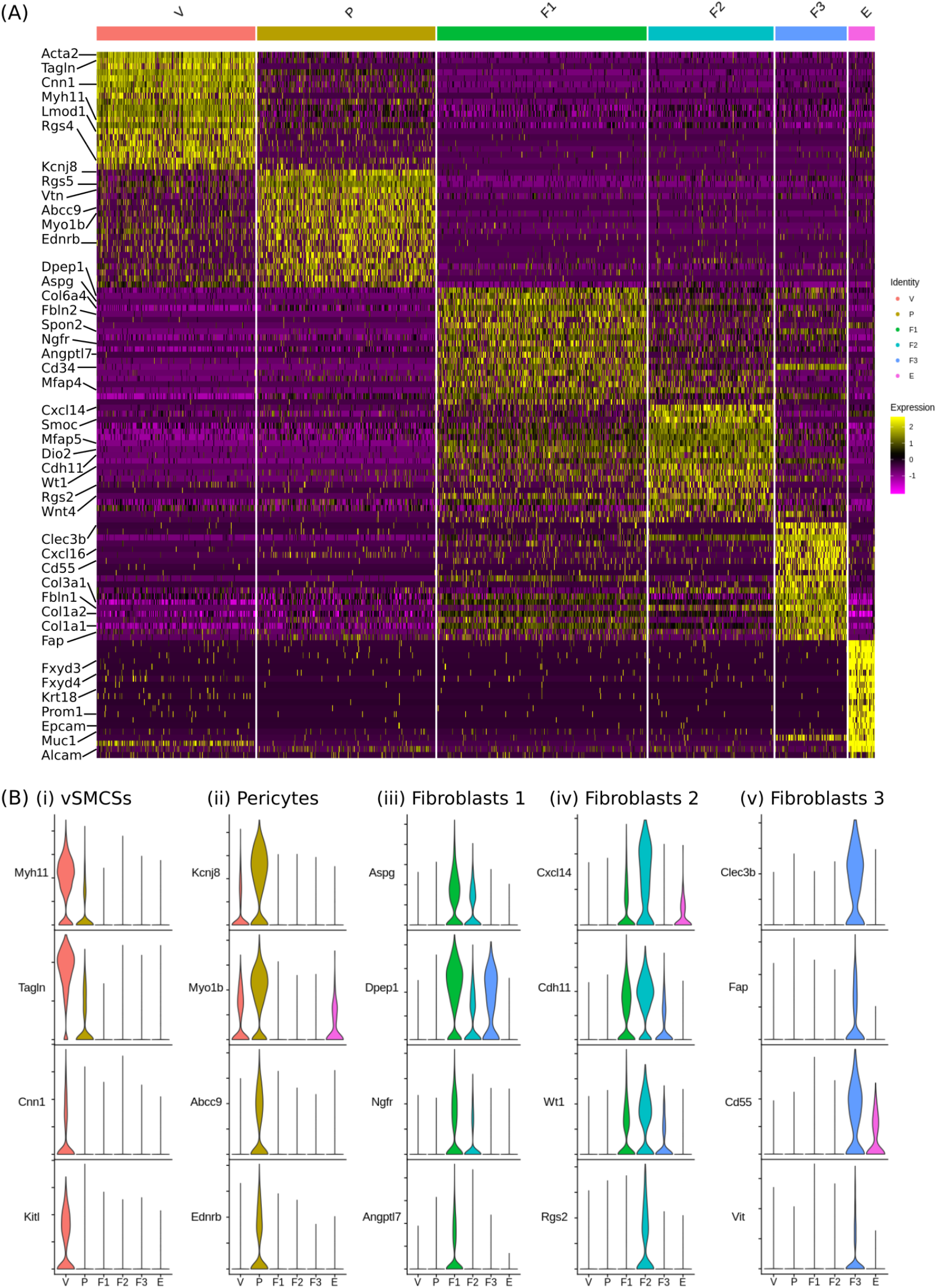
*Single-cell RNA sequencing identifies five transcriptionally discrete populations of mesenchymal cells in the GFP+ fraction of* Pdgfrb-*BAC*-eGFP mouse uterine tissue. (A) *Scaled heatmap (yellow, high; purple, low) displaying differentially expressed genes per cluster when compared to all other clusters (top is colour coded and named by cluster; V=vascular smooth muscle cells (vSMC), P=pericytes, F1= fibroblasts 1, F2= fibroblasts 2, F3= fibroblasts 3, E= epithelial cells). The expression of top twenty exemplar genes (rows) in each cell (column) is displayed. (B) Violin plot: identification of genes preferentially expressed by each cell cluster: (i)* Myh11, Tagln, Cnn1, Kitl *by vSMCs (V); (ii)* Kcnj8, Myob1, Abcc9, Ednrb *by pericytes (P); (iii)* Aspg, Dpep1, Ngfr, Angptl7 by fibroblasts 1 (F1); (iv) *Cxcl14, Cdh11*, Wt1, *Rgs2* by fibroblasts 2 (F2); *Clec3b, Fap, Cd55, Vit* by fibroblasts 3 (F3).

This analysis reveals that in the *Pdgfrb*-BAC-eGFP mouse uterus GFP^dim^ cells are represented by three subpopulations of fibroblasts while the GFP^bright^ cells are a mixture of pericytes and vSMCs.

### Gene ontology enrichment analysis reveals functional heterogeneity within the mesenchymal cell subpopulations of the endometrium during homeostasis

Differential gene expression analysis between vSMC (V) and pericyte (P) cell clusters identified genes that can be attributed to each distinct cell cluster (V: n=246, P: n=63) and those common to both (n=127) (Figure 6A-B). Gene ontology (GO) analysis revealed functional terms enriched by the transcriptomic profile of each individual cell cluster. GOBP enriched across both perivascular cell populations included circulatory system process, cell-substrate adhesion and regulation of actin filament-based process (Figure 6C). GOBP enrichment terms associated with the vSMC gene signature included muscle cell development, differentiation and contraction and the regulation of blood circulation while those associated with the pericyte gene signature included ECM organisation, smooth muscle cell migration, defence response to virus and response to interferons (Figure 6C). These results are consistent with those of the bulk mRNA sequencing study. Leading-edge analysis defined the core set of genes responsible for the enrichment of the top 10 GO terms associated with vSMCs (Figure 7A(i)) and pericytes (Figure 7A(i)). Genes that were both highly expressed and part of the leading edge/core subset for multiple GO terms were hypothesised to have biological relevance: vSMC associated genes included *Pln, Lmod1* and *Sorbs2* (Figure 7A(ii-iii)); pericyte associated genes included *Vtn, Nrp1* and *Ifitm1* (Figure 7B(ii-iii)).

**Figure 6.**
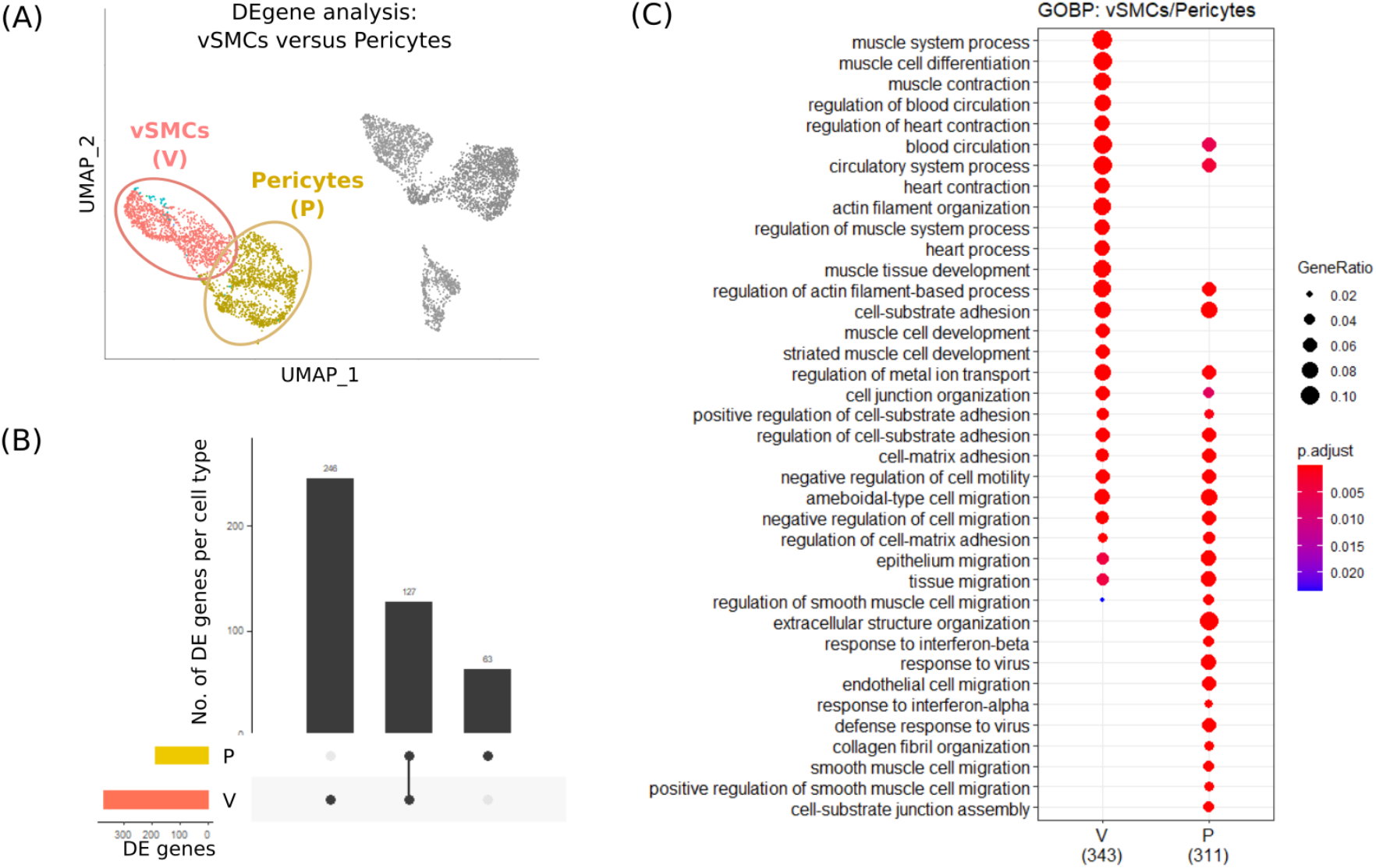
Gene ontology enrichment analysis reveals functional differences between two discrete perivascular subpopulations identified through single-cell RNA sequencing of mouse uterus. **(A)** UMAP visualisation: differential gene expression analysis carried out between vSMCs (V) and pericytes (P) to identify specific transcriptomic signatures. **(B)** Upset plot: visualisation of the intersection between gene lists for the vSMC (V) and pericyte (P) showing the number of genes that can be attributed to each cell cluster individually (V: n=246, P: n=63) and those common to both (n=127). **(C)** Dot plot: GO enrichment terms relating to biological processes (BP) associated with the signatures corresponding to the two discrete perivascular cell populations (V; P); dot size: number of genes in data/number of genes associated with GO term (gene ratio); dot colour: pvalue representing the enrichment score.

**Figure 7.**
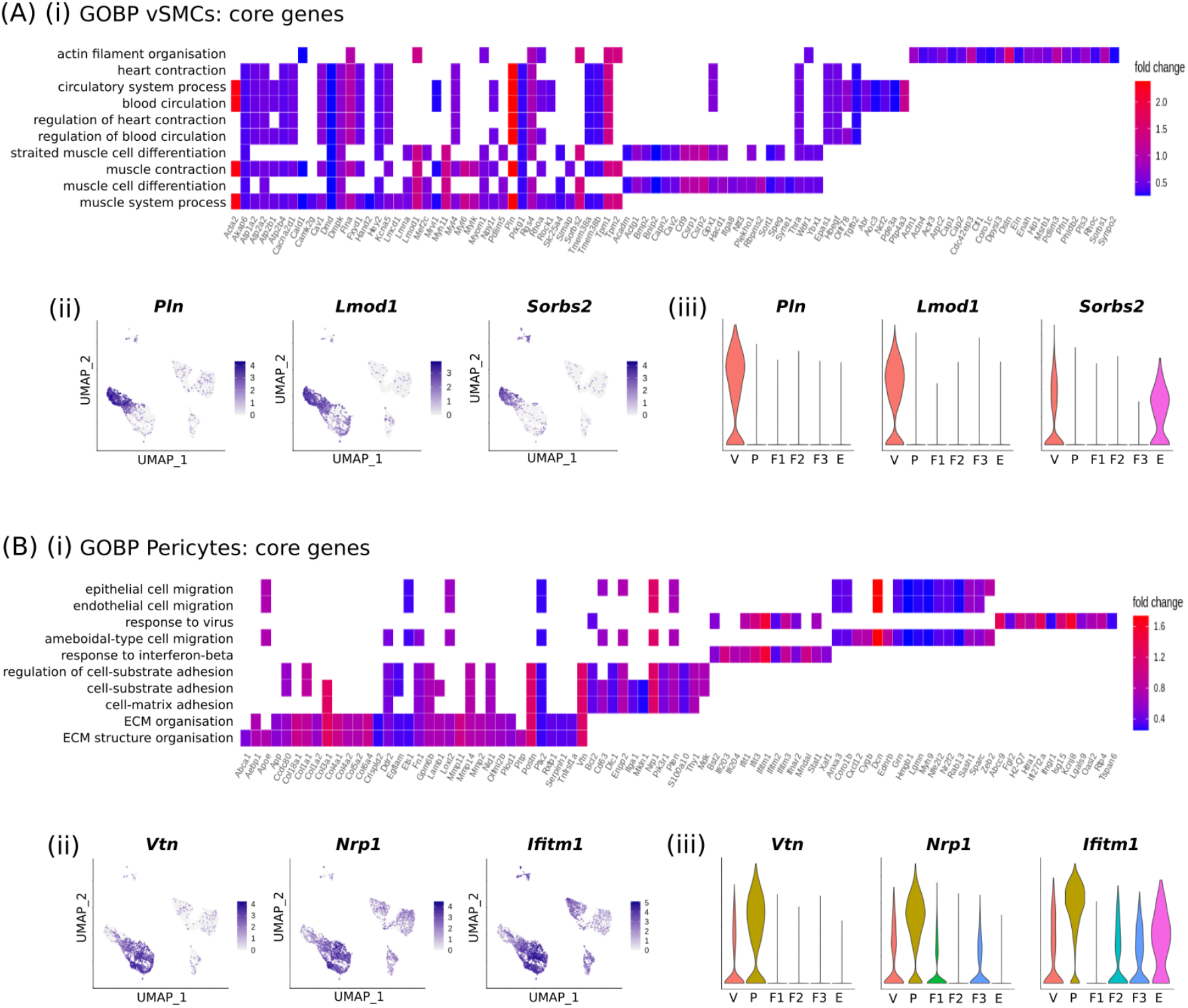
Leading-edge analysis defines the core set of genes responsible for the enrichment of the top 10 GOBP functional terms associated with perivascular cell gene signatures. **(A) (i)** Heatplot: visualisation of the overlap between core genes associated with the top 10 GOBP terms enriched for by the vSMC transcriptome; coloured by relative fold change. **(ii)** UMAP visualisation and **(iii)** Violin plot: expression of Pln, Lmod1 and Sorbs2 preferentially expressed by the vSMC cluster (V). **(B) (i)** Heatplot: visualisation of the overlap between core genes associated with the top 10 GOBP terms enriched for by the pericyte transcriptome; coloured by relative fold change. **(ii)** UMAP visualisation and **(iii)** Violin plot: expression of Vtn, Nrp1 and Ifitm1 preferentially expressed by the pericyte cluster (P).

Similarly, differential gene expression analysis between the three fibroblasts subpopulations (F1, F2, F3) revealed genes that can be attributed to each cell cluster individually (F1: n=58, F2: n=54, F3: n=132) and those common to all (n=120) or shared by various pair groups (F1&F2: n=77, F1&F3: n=48, F2&F3: n=8) (Figure 8A-B). Gene signatures that defined the fibroblast subpopulations were found to enrich for discrete GOBP functional terms: F1 enriched for the regulation of immune responses, response to interferons and antigen presentation; F2 enriched for reproductive structure development, response to wounding and tissue morphogenesis; and F3 enriched for ECM production, connective tissue development and collagen deposition (canonical fibroblast functions) (Figure 8C). Leading-edge analysis defined the core set of genes responsible for the enrichment of the top 10 GO terms associated with F1 (Figure 9A(i)), F2 (Figure 9B(i)) and F3 (Figure 9C(i)). Genes that were both highly expressed and part of the leading edge/core subset for multiple GO terms were hypothesised to have biological relevance: F1 associated genes included *Ifit1*, *Ifit3* and *H2*-*Q7* (Figure 9A(ii-iii)); F2 associated genes included *Smoc2, Wnt4* and *Aldh1a2* (Figure 9B(ii-iii)); and F3 associated genes included *Col14a1, Mmp3* and *Efemp1* (Figure 9C(ii-iii)).

**Figure 8.**
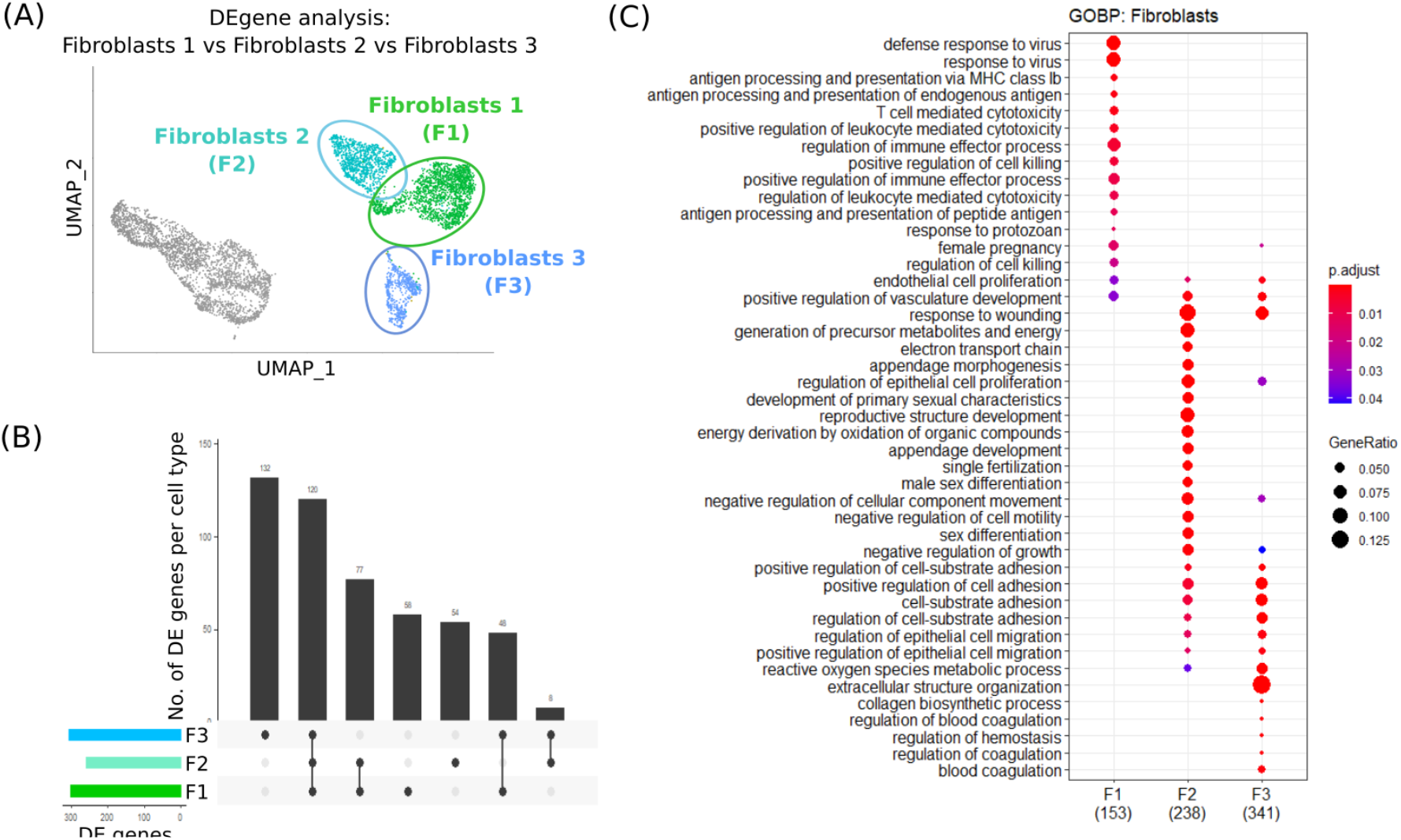
Gene ontology enrichment analysis reveals functional differences between three discrete fibroblast subpopulations identified through single-cell RNA sequencing of mouse uterus. **(A)** UMAP visualisation: differential gene expression analysis carried out between three distinct fibroblasts subpopulations (F1, F2, F3). **(B)** Upset plot: visualisation of the intersection between gene lists for the three subpopulations of stromal fibroblasts (F1, F2, F3) showing the number of genes that can be attributed to each cell cluster individually (F1: n=58, F2: n=54, F3: n=132) and those common to all (n=120) or shared by group pairs (F1&F2: n=77, F1&F3: n=48, F2&F3: n=8). **(C)** Dot plot: GO enrichment terms relating to biological processes (BP) associated with the signatures corresponding to the three discrete fibroblast populations (F1, F2, F3); dot size: number of genes in data/number of genes associated with GO term (gene ratio); dot colour: pvalue representing the enrichment score.

**Figure 9.**
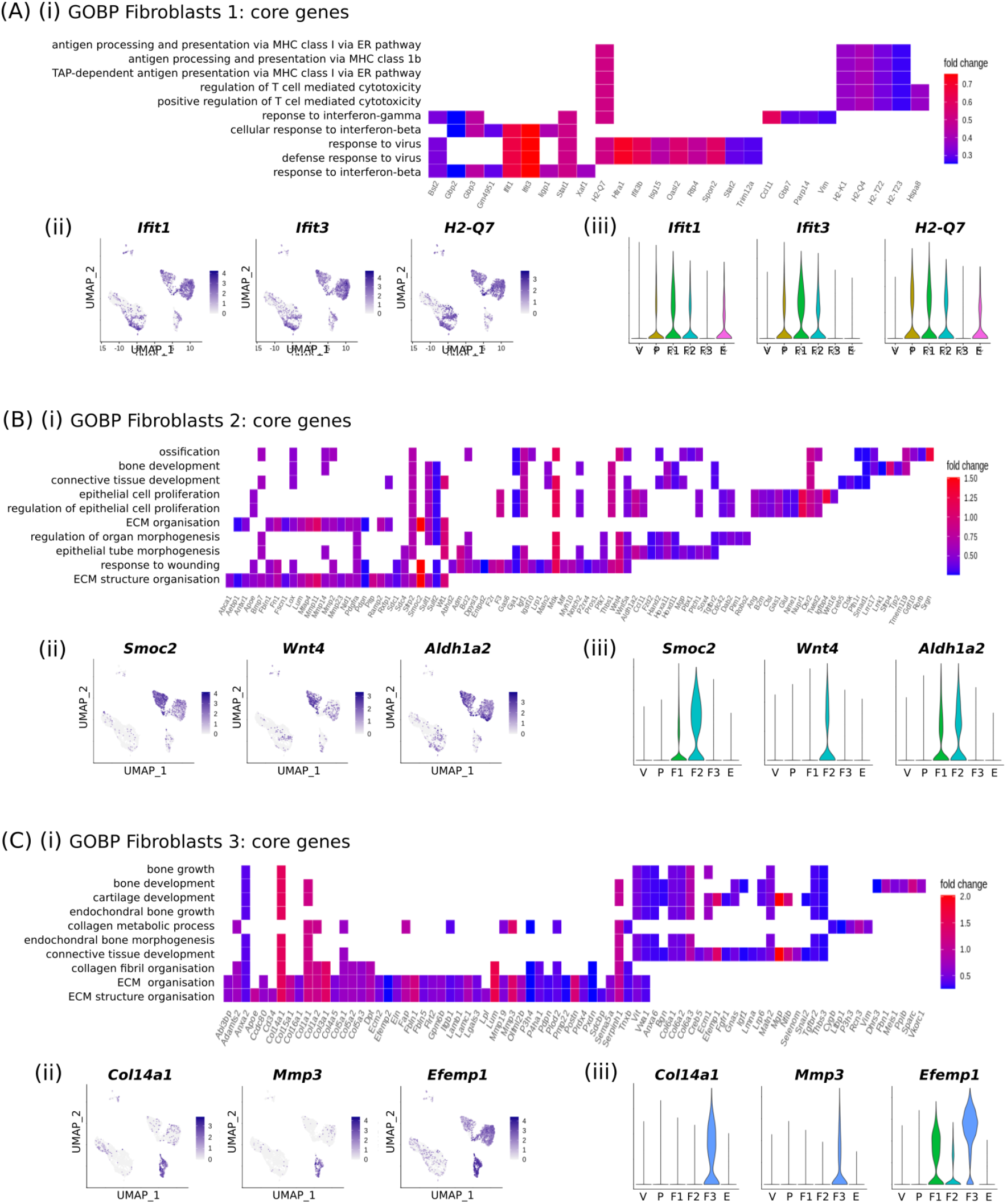
Leading-edge analysis defines the core set of genes responsible for the enrichment of the top 10 GOBP functional terms associated with fibroblasts subset gene signatures. **(A) (i)** Heatplot: visualisation of the overlap between core genes associated with the top 10 GOBP terms enriched for by the Fibroblast 1 transcriptome; coloured by relative fold change. **(ii)** UMAP visualisation and **(iii)** Violin plot: expression of Ifit1, Ifit3 and H2-Q7 preferentially expressed by the Fibroblast 1 cluster (F1). **(B) (i)** Heatplot: visualisation of the overlap between core genes associated with the top 10 GOBP terms enriched for by the Fibroblast 2 transcriptome; coloured by relative fold change. (ii) UMAP visualisation and **(iii)** Violin plot: expression of Smoc2, Wnt4 and Aldh1a2 preferentially expressed by the Fibroblast 2 cluster (F2). **(C) (i)** Heatplot: visualisation of the overlap between core genes associated with the top 10 GOBP terms enriched for by the Fibroblast 3 transcriptome; coloured by relative fold change. **(ii)** UMAP visualisation and **(iii)** Violin plot: expression of Col14a1, Mmp3 and Efemp1 preferentially expressed by the Fibroblast 3 cluster (F3).

## DISCUSSION

In this study we provide the first definitive analysis of mesenchymal cells in the adult mouse endometrium identifying five subpopulations of cells including closely related populations of pericytes and vSMC as well as three subpopulations of fibroblasts. Bioinformatics revealed that pericytes and vSMC shared functions associated with the circulatory system, actin-filament process and cell adhesion and distinct roles for subpopulations of fibroblasts in regulation of immune responses, response to wounding and organisation of extracellular matrix. These data provide the platform for comparisons between mesenchymal cells in endometrium, a tissue that exhibits remarkable resilience and regeneration, and other adult tissues such as liver which are prone to fibrosis.

In uterine tissue recovered from cycling *Pdgfrb*-BAC-eGFP transgenic mice, we established that GFP expression was specific to the stroma and identical to endogenous PDGFRβ protein. These results were consistent with the pattern of expression already described in uninjured male mouse liver which recorded co-expression of a *Pdgfrb*-driven reporter protein and endogenous PDGFRβ in hepatic stellate cells (Henderson, et al. 2013). The intensity of the GFP signal was not homogeneous, with two discrete cell populations (GFP^bright^ and GFP^dim^) easily distinguished in tissue sections. GFP^dim^CD146-were located throughout the stroma but GFP^bright^CD146+ cells were only found in close proximity to CD31+ endothelial cells. In human endometrium, stem-like progenitors have been identified in a perivascular location based on PDGFRβ and CD146 (Kaitu’u-Lino et al. 2012). We therefore conducted further studies to determine if the GFP^bright^ cells were the mouse equivalent of these human cells.

Bulk mRNA sequencing combined with bioinformatics analysis identified genes upregulated in the GFP^bright^ cells including those previously reported as being expressed by vSMCs and/or pericytes such as *Mcam* (CD146), *Acta2* (αSMA), *Myh11, Olfr78, Rgs4, Rgs5, Cspg4* (NG2), *Vcam1, Kitl* and *Kcnj8* (Cheng, et al. 2017; Gaafar, et al. 2014; He, et al. 2016; Spitzer, et al. 2012). A previous transcriptomic analysis (by gene array) of human putative endometrial mesenchymal PDGFRβ+CD146+ cells compared to stromal fibroblasts (PDGFRβ+CD146-) also identified a gene signature they reported as similar to that of pericytes (Spitzer et al. 2012). Notably, there was clear overlap between genes identified in these human cells and those we identified in the mouse GFP^bright^ cells. GOBP terms enriched by the GFP^bright^ gene signature highlighted functions such as muscle cell contraction, regulation of blood circulation and regulation of leukocyte migration, consistent with functions previously ascribed to perivascular smooth muscle cells and/or pericytes.

Validation experiments using purified cells revealed that putative perivascular cell mRNAs including *Mcam, Vcam1* and *Kitl* were also expressed by CD31+ endothelial cells. In contrast, expression of *Acta2* (αSMA), *Myh11, Olfr78, Kcnj8, Cspg4* (NG2), *Rgs4* and *Rgs5* appeared to be specific to GFP^bright^ cells and importantly not detectable in either stromal (GFP^dim^CD146-) or endothelial (CD31+) cells. Immunoanalysis of the pericyte/vSMC-specific proteins Cspg4 (NG2), αSMA (Acta2) and MYH11 (Ansell and Izeta 2015; Crisan, et al. 2008; Mills, et al. 2013) found that NG2 was expressed by GFP^bright^ cells throughout the endometrium whereas αSMA and MYH11 were only detected in a subset of the GFP^bright^ cells located at the endometrial/myometrial junction. We propose NG2 (*Cspg4*) should be used to identify perivascular cells in mouse endometrium in preference to CD146 (*Mcam*) alone or in combination with PDGFRβ+. These findings also call into question interpretation of results in previous genomic studies which attributed a mixture of vSMC/pericyte genes and associated functions to putative endometrial pericytes (Guimaraes-Camboa, et al. 2017; Spitzer et al. 2012).

Tissue analysis of CD31+ and Cspg4/NG2+ cells revealed that every NG2+ cell was colocated with a CD31+ endothelial cell consistent with them occupying a perivascular niche. The ratio of NG2:CD31 expressing cells was calculated to be 1.25:1 (±0.02). In the literature, pericyte coverage of blood vessels is described as dense or sparse and is thought to be related to the extent of substance transfer between blood vessels in different tissues. For example, the retina has a high ratio of 1:1 while the peripheral vasculature has a low ratio of 1:100 (Andersson, et al. 2015). A higher ratio is believed to be related to greater regulation of blood flow and more frequent remodelling of micro-vessels (Andersson et al. 2015). Coculture studies have shown that a 1:1 pericyte:endothelial cell ratio significantly inhibits endothelial cell proliferation when compared to a ratio of 1:10 or 1:20, believed to be due to an inability of pericytes to communicate with the increasing number of endothelial cells (Bodnar, et al. 2016). Our result suggest endometrial vasculature in the mouse has a dense coverage of perivascular cells which may suggest they play a role in stabilisation of vessels or limiting transfer from the blood.

To further understand and dissect cellular heterogeneity within the endometrial mesenchyme we performed single cell RNA sequencing on the total population of GFP+ endometrial cells. Unsupervised cluster analysis of the resulting dataset identified six distinct clusters, five of which were classified as mesenchyme (GFP+/*Pdgfrb+*) with a small sixth cluster representing contaminating *Pdgfrb*-epithelial cells. Of the mesenchyme clusters, two closely related clusters were *Pdgfrb^high^/Mcam+*, equivalent to GFP^bright^ cells, while three distinct clusters of *Pdgfrb^low^/Mcam*- (GFP^dim^ equivalent) cells were also observed.

The unexpected heterogeneity within the *Pdgfrb^low^/Mcam*-cells, consistent with the existence of three fibroblasts subpopulations (F1, F2, F3), prompted further analysis as human endometrial stromal fibroblasts (eSFs) are usually considered a homogenous population of cells with a shared set of functions in endometrial physiology. A role for human eSFs in modulating the immune response has been described using an *in vitro* co-culture system (Queckborner, et al. 2020). More recently, using an elegant multi-omics profiling and integrative bioinformatics approach, an important role for so called structural cells (endothelial, epithelial cells; fibroblasts) in immune cell regulation has been confirmed in 12 organs of the mouse, during both homeostasis and in response to immunological challenges (Krausgruber, et al. 2020). Notably, although this study did not include reproductive tissues such as the uterus, many of the identified immune genes and associated GOBP terms are reflected in our scRNAseq results, primarily enriched in F1 (defence response to virus, cellular response to IFNg, antigen processing and presentation). Our results further suggest that such roles can be attributed to specific subpopulations of cells, something that is not accounted for by methods employed by Krausgruber *et al*. However, both reports highlight that structural cell populations may have previously underappreciated roles in tissue homeostasis out with the typical barrier/connective tissue functions (Krausgruber et al. 2020).

Comparative single cell analysis of stromal cells from primary cultures and biopsies of human endometrium has identified many differences suggesting cultured cells may not represent the full range of *in vivo* phenotypes (Krjutskov, et al. 2016). This potentially undermines the relevance of results obtained using cultured cells alone. ScRNAseq has been used to investigate changes in stromal cell fibroblasts during the process of decidualization in women with time-dependent changes in gene signatures identified in fibroblasts as they transformed into decidual cells (Lucas, et al. 2020). However, scRNAseq performed on ~3000 cells from an endometrial biopsy defined only a single ‘fibroblast’ cluster which had a genetic signature implicated in angiogenesis and wound healing. This study noted that stromal fibroblasts upregulated immunomodulatory genes during decidualization including IL15, consistent with previous reports that fibroblast/decidual cells-derived factors play a key role in immune cell recruitment (Kitaya, et al. 2005). Heterogeneity within the stromal fibroblast population of the human biopsies was not reported probably because too few cells were present in the pooled cell samples. In a paper published on Bioarchiv [https://www.biorxiv.org/content/10.1101/350538v2.full] that used 2149 single cells from 19 healthy human endometria across the menstrual cycle likewise all fibroblasts were assigned to a single cell cluster. In contrast in the current study we were able to target our analysis to a purified population of 5000 GFP+ cells detecting 18,000 unique genes. Further work with larger numbers of stromal cells from human tissues will be necessary to determine whether cell populations equivalent to F1, F2 and F3 exist in the human endometrium.

In recent years there has been a massive increase in the range of tissues from mouse and other species analysed using scRNAseq technologies. Many datasets are available, including those hosted by EMBL-EBI (Papatheodorou, et al. 2020), the single cell expression atlas: [https://www.ebi.ac.uk/gxa/sc/home], (Tabula Muris, et al. 2018) and a Single Cell Atlas generated by Han et al (Han, et al. 2018); https://www.ncbi.nlm.nih.gov/geo/query/acc.cgi?acc=GSE108097). Unfortunately, the Tabula Muris did not include any reproductive tissues in the 20 reported in their 2018 publication (Tabula Muris, Overall et al. 2018). However Han and colleagues reported single cell analysis of 3761 cells from the mouse uterus with identification of 19 cell clusters. Cross-comparison between stromal cells and tissue-resident macrophages isolated from uterus and other tissues suggested they had a unique and distinct signature but no further validation was provided (see figure 6 in (Han et al. 2018).

We believe the data described in our study is the first comprehensive analysis of mouse endometrial mesenchyme in the adult mouse generated using scRNAseq. A study by Saatcioglu and colleagues used similar techniques to investigate gene signatures in the immature mouse uterus on post-natal day 6 of both controls and those treated with Mullerian Inhibiting Substance (MIS/AMH; (Saatcioglu, et al. 2019). The authors identified two closely related populations of stromal cells which they designated as ‘inner’ and ‘outer’ and a small population of ‘pericytes’. Although the tissue used was immature and total uterine tissue was analysed there is notable overlap between stromal candidate gene markers we discovered and those in their dataset, including *Smoc2, Cdh11, Col6a4* (Saatcioglu et al. 2019) reinforcing our data analysis strategy.

## Conclusion

The current study has used immunohistochemistry, flow cytometry, qRTPCR and sequencing methods to shed new light on heterogeneity within mesenchymal cells in mouse endometrium. We have demonstrated that although PDGFRβ and CD146 are reported to be ubiquitously expressed by pericytes in a wide range of tissues (Bodnar et al. 2016) in endometrium they are not exclusive to this cell type. We propose that NG2 (*Cspg4*) is a more specific and exclusive marker of endometrial cells residing in the perivascular niche (vSMC/pericytes) in mice.

Our scRNAseq analysis revealed novel heterogeneity in the endometrial mesenchyme (Figure 10). We believe these data provide a platform for comparison between single cell data from endometrium, a tissue that exhibits remarkable resilience and is not prone to fibrotic transformation and other tissues such as liver (Dobie, et al. 2019; Ramachandran, et al. 2020), lung (Peyser, et al. 2019) and kidney (Kirita, et al. 2019; Kirita, et al. 2020), in which the role of fibroblasts in fibrotic responses has been studied. Importantly they also enhance our capacity for *in vivo* targeting and lineage tracing of defined subpopulations of cells within the endometrial stroma to inform our understanding of their role(s) in endometrial function.

**Figure 10.**
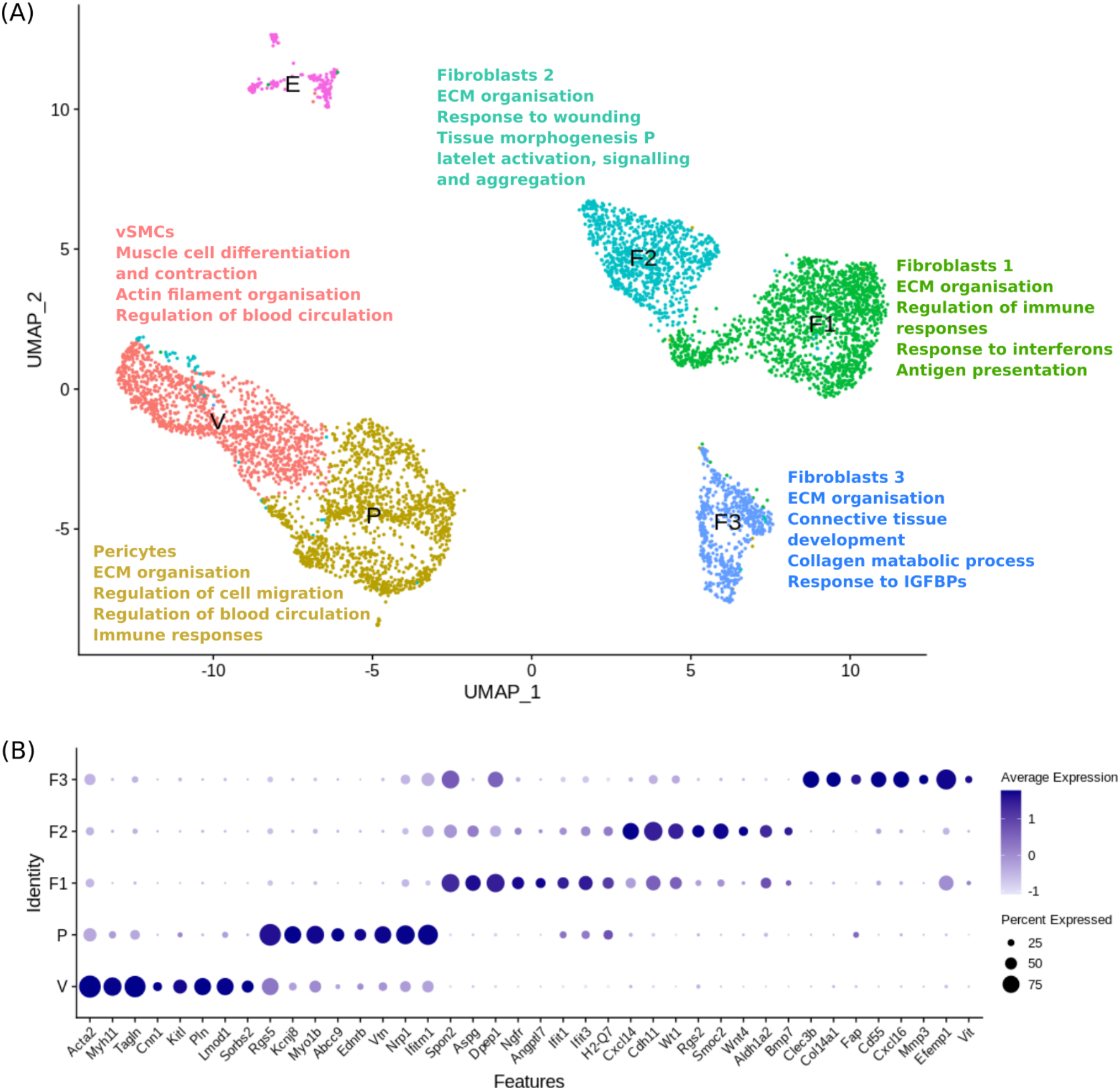
Single cell sequencing analysis identifies five transcriptomically and functionally discrete subpopulations of PDGFRβ+ mesenchymal cells in the mouse endometrium. **(A)** UMAP visualisation: gene signatures associated with each individual cell cluster enrich for distinct biological functions. **(B)** Dotplot: exemplar genes identified through phenotypic and functional analyses of transcriptomic signatures that are preferentially expressed by each individual cell cluster (vSMCs (V): Acta2, Myh11, Tagln, Cnn1, Kitl, Pln, Lmod1, Sorbs2; pericytes (P): Rgs5, kcnj8, Myo1b, Abcc9, Ednrb, Vtn, Nrp1, Ifitm1; fibroblasts 1 (F1): Spon2, Aspg, Dpepe1, Ngfr, Angptl7, Ifit1, Ifit3, H2-Q7; fibroblasts 2 (F2): Cxcl14, Cdh11, Wt1, Rgs2, Smoc2, Wnt4, Aldh1a2, Bmp7; fibroblasts 3 (F3): Clec3b, Col14a1, Fap, Cd55, Cxcl16, Mmp3, Efemp1, Vit; epithelial cells (E): Cdh1, Epcam, Krt7, Krt18, Muc1, Muc4, Prom1, Alcam. Dot size: percentage of cells expressing the gene; dot colour: average expression of gene.

## MATERIALS AND METHODS

### Animals

In *Pdgfrb*-BAC-eGFP reporter mice (C57BL/6 background), eGFP expression is driven by the regulatory sequences of the *Pdgfrb* gene and is therefore expressed by all cells in which this promoter is active. The use of this transgenic mouse for studies on liver mesenchymal cells has previously been described (Henderson et al. 2013): these mice were originally obtained from GENSAT and deposited in MMRRC-STOCK Tg(Pdgfrb-EGFP)JN169Gsat/Mmucd, 031796-UCD). All mice were genotyped at weaning as described previously (Henderson et al. 2013). Uterine tissue samples were collected from cycling adult female mice (8-10 weeks old) with oestrous stage confirmed by vaginal smears as described in (Caligioni 2009).

### Tissue fixation

Uterine tissue samples were fixed in 4% (w/v) PFA for 2h at 4°C, rinsed thoroughly in PBS and stored overnight in 18% (w/v) sucrose at 4°C. Approximately 24h later samples were embedded in OCT medium and stored at −80°C. Frozen tissue sections (5μm) were cut and mounted onto Xtra adhesive pre-cleaned micro slides (Surgipath, Leica Biosystems) and airdried at room temperature for 30 minutes prior to staining (minimum of 2 sections of each uterine horn per mouse).

### Immunohistochemistry

Haematoxylin and Eosin (H&E) staining was performed according to standard methods. For immunofluorescence tissue sections were washed in PBS to remove residual OCT and incubated with 3% (v/v) hydrogen peroxide solution in methanol for 30 min at room temperature, washed (unless stated otherwise, all wash steps included one 5 min wash in PBS containing 0.05% (v/v) Tween20 and two 5 min washes in PBS) and further incubated with 20% (v/v) normal goat serum (NGS: PBS containing 20% goat serum and 0.05% (w/v) bovine serum albumin (BSA)) for 30 min at room temperature. Sections were washed and incubated overnight at 4°C with the primary antibody at an optimised dilution in NGS. Following a further wash, sections were incubated with an HRP-conjugated secondary antibody at an optimised dilution in NGS for 30 min at room temperature followed by a 10 min incubation with Tyramide solution (PerkinElmer). Primary and secondary antibodies and associated working dilutions are given in Supplementary Table 1. Co-stains were completed sequentially. Sections were counterstained with DAPI (Sigma, D9542), overlaid with Permaflour (Immunotech) and mounted with coverslips (VWR Prolabo). Images were captured on a Zeiss LSM 510 Meta Confocal microscope using Zen 2009 software (Zeiss).

### Flow cytometry and fluorescence-activated cell sorting

Tissue processing for flow cytometry analysis and FACS was performed as previously described (Cousins et al. 2016). Cells were incubated for 30 min on ice with optimised dilutions of fluorescently-conjugated antibodies as detailed in Supplementary Table 2. Cell pellets re-suspended in 500μl PBS at 4°C were analysed using a BD 5L LSR Fortessa and BD FACSDiva software (BD Biosciences). To exclude dead cells, DAPI (1/10K) was added prior to flow cytometry analysis. Cells were sorted using a FACS Aria II instrument and BD FACSDiva software (BD Biosciences). Data analysis was performed using FlowJo analysis software (FlowJo LLC, Oregon, USA).

### RNA and cDNA preparation for quantitative real-time PCR

For RNA extraction of tissue, samples were added to TriReagent (Sigma, Cat. No. T9424) and processed as detailed in (Cousins et al. 2014). The total concentration and purity of resulting RNA was determined using a Nanodrop ND100 Spectrophotometer (Thermo Fisher Scientific). Reverse transcription of RNA to cDNA was performed with the Superscript VILOTM cDNA synthesis kit (Invitrogen, Cat. No. 11754-250) according to the manufacturer’s instructions and the following PCR settings: 25°C for 10 min, 42°C for 60 min and 85°C for 5 min in a Peltier Thermal Cycler (PTC-200).

RNA extraction of FACS-isolated cells was carried out using the automated Maxwell^®^ Instrument (Promega, Cat. No. AS2000, Wisconsin, USA). The SimplyRNA cells protocol was run using the Maxwell^®^ 16 Instrument configured with the Maxwell^®^ 16 High Strength LEV Magnetic Rod and Plunger Bar Adaptor (Promega, Cat. No. SP10790) and Maxwell^®^ 16 firmware version 4.95. The total concentration and purity of resulting RNA was determined using the RNA Pico Sensitivity Assay (Perkin Elmer, Cat. No. CLS960012) as per manufacturer’s instructions and loaded onto the DNA 5K/RNA/Charge Variant Assay LabChip (Perkin Elmer, Cat. No. 760435) to be read using a LabChip GX Touch Nucleic Acid Analyser (Perkin Elmer, Cat. No. CLS138162). Reverse transcription and amplification of RNA purified from isolated cells was performed with the NuGEN Ovation RNA-Seq System V2 (NuGEN Cat. No. 7102-32) according to manufacturer’s instructions. The integrity of resultant cDNA was assessed using a LabChip GX Touch 24 Nucleic Acid Analyzer as per manufacturer’s instructions (Perkin Elmer).

### Quantitative RT-PCR

The Applied Biosystems TaqMan method (Thermo Fisher Scientific) was used to detect specific PCR products. Primers for genes of interest were designed by the Universal Probe Library Assay Design Centre (Roche Applied Science, Penzberg, Germany) and purchased from Eurofins MWG Operon (Ebersberg, Germany) (sequences in Supplementary Table 3). Reactions were prepared in duplicate and amplification performed at 95°C for 5 min and then 30 cycles of 95°C for 1 min, 58°C for 1 min, and 72°C for 1 min on a real time PCR system (Quantstudio5). Relative expression for each gene was calculated using the standard curve method in which the amount of target genes was normalised to beta-actin (*Actb*) and relative expression between samples was calculated. Normalised expression values are displayed as a fold-change in expression. Statistical analyses were performed using Graphpad Prism software. In data that was normally distributed a student’s t-test was performed to determine the significance of a difference between two groups. When comparing the means of more than two groups a one-way ANOVA was used followed by a multiple comparisons test such as Sidak’s or Tukey’s. All data is presented as mean±SEM and criteria for significance is p<0.05.

### Illumina based mRNA sequencing

Endometrial mesenchymal cells (GFP+) were isolated from cycling *Pdgfrb*-BAC-eGFP mice by FACS and RNA extracted using the Maxwell^®^ SimplyRNA cells kit (Promega, Cat. No. AS1270). cDNA was amplified using the NuGEN Ovation V2 system and all purified samples were judged to be of high quality and sufficient quantity for next generation mRNA sequencing using the LabChip method (Perkin Elmer). TruSeq DNA Nano gel free libraries (350bp insert) were prepared for each sample and high-throughput sequencing of library products was performed on the HiSeq 4000 75PE platform according to the standard protocols by Edinburgh Genomics (http://genomics.ed.ac.uk).

### Bioinformatics analysis of mRNA sequencing

FASTQ files were used for genome alignment: Cutadapt was used to filter poor quality raw reads (threshold 30) and trim adapter sequences of the NuGen Ovation V2 Kit (AGATCGGAAGAGC). After trimming, reads were filtered to have a minimum length of 50bases then aligned to the Mus musculus genome from Ensembl (assembly GRCm38, annotation version 84) using STAR2 version 2.5.2b. Reads were assigned to exon features and grouped by gene ID in the reference genome using featurecounts3. The raw counts table was filtered to remove rows consisting predominantly of near-zero counts, filtering on counts per million (CPM) to avoid artefacts due to library depth. Principal components analysis was undertaken on normalised and filtered expression data to explore observed patterns with respect to experimental factors. The cumulative proportion of variance associated with each factor was used to study the level of structure in the data, while associations between continuous value ranges in principlal components and categorical factors were assessed with an ANOVA test. Points were assigned as outliers in each component if they occurred outside the interquartile range +1.5. If appropriate, problematic samples were excluded from downstream analysis. Differential analysis was carried out using EdgeR4 (version 3.16.5; (Robinson, et al. 2010) and a quasi-likelihood (QL) F-test performed using the desired contrasts. Gene set enrichment analysis of differentially expressed gene sets was carried out using the ‘clusterProfiler’ R package (Yu, et al. 2012) with R version 4.0.2.

Ten candidate genes were chosen based on their CPM and fold change between experimental groups for validation studies: expression was analysed by qPCR using a new set of cell samples generated from *Pdgfrb*-BAC-eGFP mouse endometrial tissues. Fold changes were calculated by comparing relative expression values to that of the housekeeper gene *Actb*.

### Chromium single cell gene expression analysis-10x Genomics

Endometrial mesenchymal cells (GFP+) were isolated from cycling *Pdgfrb*-BAC-eGFP mice by FACS previously described. Samples were pooled from uteri of 4 mice (estrus phase as determined by vaginal smearing, 2 horns pooled from each mouse: 25,000 GFP+ cells/sample giving a total number of 100,000 GFP+ cells for downstream application). Following the sort, isolated cell suspensions were counted, and viability confirmed to be >85% using a TC20™ Automated Cell Counter (BioRad, Cat. No. 1450102). Cells were partitioned into nanolitre-scale Gel bead-in-Emulsions (GEMs) containing unique 10x barcodes using the 10x Chromium™ Controller (10x Genomics, USA). cDNA libraries were generated and amplified using the Chromium™ Single Cell 3’ Library & Gel Bead Kit V2 (10x Genomics, Cat. No. 120267) and the Chromium™ Single Cell A Chip Kit 16 (10x Genomics, Cat. No. 1000009) following manufacturer’s instructions. cDNA concentration and quality were measured using the LabChip GX Touch Nucleic Acid Analyser and confirmed to surpass threshold values. Sequence data was generated on the Illumina NovaSeq platform using bespoke 10x parameters, according to the standard protocols at the facility (Edinburgh Genomics: http://genomics.ed.ac.uk/, Edinburgh).

### Bioinformatics analysis of single cell mRNA sequencing

Pre-processing of raw sequencing data files was performed using 10x Cell Ranger (version 2.0.1; https://www.10xgenomics.com). The ‘cellranger_mkfastq’ command was used to demultiplex raw base call (BCL) files generated by the Illumina sequencer, specifying the SI-GA indices associated with each sample to map individual reads back to the individual input cells. Resultant FASTQ files for each sample were fed into ‘cellranger_count’ along with the transcriptome ‘refdata-cellranger-mm10-1.2.0’ as supplied by 10x genomics, to perform genome alignment, filtering, barcode counting and UMI counting. Sequence saturation was 85.1% and 85.6% reads mapped confidently to the genome (Figure S5A). For downstream QC, clustering and gene expression analysis the *seurat* R package (V3) was utilised (Satija, et al. 2015) with R version 4.0.2. The ‘Read10x’ function was used to read in the output of the Cell Ranger pipeline returning a unique molecular identified (UMI) count matrix representing the number of molecules for each gene detected in each cell. The ‘CreateSeuratObject’ command was used to create the Seurat object used in all subsequent analyses.

The standard Seurat pre-processing workflow was followed to filter cells based on QC metrics, normalise and scale the data and finally detect highly variable features in the data. Before analysing the single-cell gene expression data we ensured that all cellular barcode data corresponded to viable cells by assessing four QC covariates: number of unique genes detected in each cell (nFeature_RNA >200 & <5000), the total number of molecules detected within a cell (nCount_RNA), the percentage of reads that map to the mitochondrial genome (<5%) and the percentage of reads that map to ribosomal proteins (<5%). These QC covariates were used to identify non-viable cells or doublets which were filtered out by thresholding prior to downstream analysis (Figure S5B-E). Resulting data was normalised by employing a global-scaling normalization method ‘LogNormalize’ that normalises the feature expression measurements for each cell by the total expression, multiplies this by a scale factor and log-transforms the result. A linear transformation (scaling) was performed to give equal weight to genes in downstream analyses so that highly expressed genes did not dominate, before a principal component analysis was performed as the chosen linear dimensional reduction method. Unsupervised clustering based on the first 20 principal components of the most variably expressed genes was performed using a graph based approach (‘FindNeighbours’, ‘FindClusters’; resolution= 0.2) which embeds cells in a K-nearest neighbour (KNN) graph based on the euclidean distance in PCA space with edges drawn between cells with similar feature expression patterns which is then partitioned into highly interconnected ‘quasi-cliques’ or ‘communities’. Resultant clusters were visualised using the manifold approximation and projection (UMAP) method.

Differential gene expression analysis was performed (‘FindAllMarkers’) to identify genes expressed by each cell cluster when compared to all other clusters, based on the nonparametric Wilcoxon rank sum test. Outputs were visualised using the ‘DoHeatmap’, ‘FeaturePlot’ and ‘VlnPlot’ functions. Cluster identification was determined by analysing the expression of canonical cell markers across the cell clusters. The ‘UpsetR’ package (Conway, et al. 2017) was used to analyse the co-expression of upregulated genes across cell clusters and compare differentially expressed gene lists generated by NGS and SCseq methods. Over-represented functional annotations in the differentially expressed genes were detected by using the ‘clusterProfiler’ package (Yu et al. 2012) using core functions to interpret data in the context of biological function, pathways and networks.

## Supporting information

Main manuscript

## Acknowledgements

Funding for this research was provided by a MRC PhD Studentship in Tissue Repair and a MRC Transition Fellowship to PWK funded by a Doctoral Training Award to the University of Edinburgh (MR/N013166/1); DAG and AZ-E were funded by a MRC Programme Grant to PTKS (MR/N024524/1); JRS was supported by a Wellcome Trust funded Edinburgh Clinical Academic Track (ECAT) Fellowship; NCH was supported by a Wellcome Trust Senior Research Fellowship in Clinical Science (219542/Z/19/Z). We thank staff supporting core facilities including flow cytometry and animal husbandry for their support and guidance. We are grateful to Beth Henderson for her expert technical help in setting up the 10X sequencing and to Dr Isaac Shaw for his insights and editorial comments.

## REFERENCES

Andersson E, Zetterberg E, Vedin I, Hultenby K, Palmblad J & Mints M 2015 Low pericyte coverage of endometrial microvessels in heavy menstrual bleeding correlates with the microvessel expression of VEGF-A. Int J Mol Med 35 433–438.

Ansell DM & Izeta A 2015 Pericytes in wound healing: friend or foe? Exp Dermatol 24 833–834.

Bellofiore N, Cousins F, Temple-Smith P, Dickinson H & Evans J 2018a A missing piece: the spiny mouse and the puzzle of menstruating species. J Mol Endocrinol 61 R25–R41.

Bellofiore N, Rana S, Dickinson H, Temple-Smith P & Evans J 2018b Characterization of human-like menstruation in the spiny mouse: comparative studies with the human and induced mouse model. Hum Reprod 33 1715–1726.

Bodnar RJ, Satish L, Yates CC & Wells A 2016 Pericytes: A newly recognized player in wound healing. Wound Repair Regen 24 204–214.

Caligioni CS 2009 Assessing reproductive status/stages in mice. Curr Protoc Neurosci Appendix 4 Appendix 4I.

Cervello I, Martinez-Conejero JA, Horcajadas JA, Pellicer A & Simon C 2007 Identification, characterization and co-localization of label-retaining cell population in mouse endometrium with typical undifferentiated markers. Hum Reprod 22 45–51.

Chan RW & Gargett CE 2006 Identification of Label Retaining Cells in Mouse Endometrium. Stem Cells.

Chan RW, Schwab KE & Gargett CE 2004 Clonogenicity of human endometrial epithelial and stromal cells. Biol Reprod 70 1738–1750.

Cheng Y, Li L, Wang D, Guo Q, He Y, Liang T, Sun L, Wang X, Cheng Y & Zhang G 2017 Characteristics of Human Endometrium-Derived Mesenchymal Stem Cells and Their Tropism to Endometriosis. Stem Cells Int 2017 4794827.

Conway JR, Lex A & Gehlenborg N 2017 UpSetR: an R package for the visualization of intersecting sets and their properties. Bioinformatics 33 2938–2940.

Cousins FL, Kirkwood PM, Saunders PT & Gibson DA 2016 Evidence for a dynamic role for mononuclear phagocytes during endometrial repair and remodelling. Sci Rep 6 36748.

Cousins FL, Murray A, Esnal A, Gibson DA, Critchley HO & Saunders PT 2014 Evidence from a mouse model that epithelial cell migration and mesenchymal-epithelial transition contribute to rapid restoration of uterine tissue integrity during menstruation. PLoS One 9 e86378.

Crisan M, Yap S, Casteilla L, Chen CW, Corselli M, Park TS, Andriolo G, Sun B, Zheng B, Zhang L, et al. 2008 A perivascular origin for mesenchymal stem cells in multiple human organs. Cell Stem Cell 3 301–313.

Dobie R, Wilson-Kanamori JR, Henderson BEP, Smith JR, Matchett KP, Portman JR, Wallenborg K, Picelli S, Zagorska A, Pendem SV, et al. 2019 Single-Cell Transcriptomics Uncovers Zonation of Function in the Mesenchyme during Liver Fibrosis. Cell Rep 29 1832–1847 e1838.

Gaafar T, Osman O, Osman A, Attia W, Hamza H & El Hawary R 2014 Gene expression profiling of endometrium versus bone marrow-derived mesenchymal stem cells: upregulation of cytokine genes. Mol Cell Biochem 395 29–43.

Gargett CE & Masuda H 2010 Adult stem cells in the endometrium. Mol Hum Reprod 16 818–834.

Gargett CE, Schwab KE & Deane JA 2016 Endometrial stem/progenitor cells: the first 10 years. Hum Reprod Update 22 137–163.

Gargett CE, Schwab KE, Zillwood RM, Nguyen HP & Wu D 2009 Isolation and culture of epithelial progenitors and mesenchymal stem cells from human endometrium. Biol Reprod 80 1136–1145.

Garry R, Hart R, Karthigasu KA & Burke C 2009 A re-appraisal of the morphological changes within the endometrium during menstruation: a hysteroscopic, histological and scanning electron microscopic study. Hum Reprod 24 1393–1401.

Gellersen B, Brosens IA & Brosens JJ 2007 Decidualization of the human endometrium: mechanisms, functions, and clinical perspectives. Semin Reprod Med 25 445–453.

Guimaraes-Camboa N, Cattaneo P, Sun Y, Moore-Morris T, Gu Y, Dalton ND, Rockenstein E, Masliah E, Peterson KL, Stallcup WB, et al. 2017 Pericytes of Multiple Organs Do Not Behave as Mesenchymal Stem Cells In Vivo. Cell Stem Cell 20 345–359 e345.

Han X, Wang R, Zhou Y, Fei L, Sun H, Lai S, Saadatpour A, Zhou Z, Chen H, Ye F, et al. 2018 Mapping the Mouse Cell Atlas by Microwell-Seq. Cell 173 1307.

He L, Vanlandewijck M, Raschperger E, Andaloussi Mae M, Jung B, Lebouvier T, Ando K, Hofmann J, Keller A & Betsholtz C 2016 Analysis of the brain mural cell transcriptome. Sci Rep 6 35108.

Henderson NC, Arnold TD, Katamura Y, Giacomini MM, Rodriguez JD, McCarty JH, Pellicoro A, Raschperger E, Betsholtz C, Ruminski PG, et al. 2013 Targeting of alphav integrin identifies a core molecular pathway that regulates fibrosis in several organs. Nat Med 19 1617–1624.

Kaitu’u-Lino TJ, Ye L, Salamonsen LA, Girling JE & Gargett CE 2012 Identification of label-retaining perivascular cells in a mouse model of endometrial decidualization, breakdown, and repair. Biol Reprod 86 184.

Kirita Y, Chang-Panesso M & Humphreys BD 2019 Recent Insights into Kidney Injury and Repair from Transcriptomic Analyses. Nephron 143 162–165.

Kirita Y, Wu H, Uchimura K, Wilson PC & Humphreys BD 2020 Cell profiling of mouse acute kidney injury reveals conserved cellular responses to injury. Proc Natl Acad Sci U S A.

Kitaya K, Yamaguchi T & Honjo H 2005 Central role of interleukin-15 in postovulatory recruitment of peripheral blood CD16(-) natural killer cells into human endometrium. J Clin Endocrinol Metab 90 2932–2940.

Krausgruber T, Fortelny N, Fife-Gernedl V, Senekowitsch M, Schuster LC, Lercher A, Nemc A, Schmidl C, Rendeiro AF, Bergthaler A, et al. 2020 Structural cells are key regulators of organ-specific immune responses. Nature 583 296–302.

Krjutskov K, Katayama S, Saare M, Vera-Rodriguez M, Lubenets D, Samuel K, Laisk-Podar T, Teder H, Einarsdottir E, Salumets A, et al. 2016 Single-cell transcriptome analysis of endometrial tissue. Hum Reprod 31 844–853.

Letouzey V, Tan KS, Deane JA, Ulrich D, Gurung S, Ong YR & Gargett CE 2015 Isolation and characterisation of mesenchymal stem/stromal cells in the ovine endometrium. PLoS One 10 e0127531.

Lucas ES, Vrljicak P, Muter J, Diniz-da-Costa MM, Brighton PJ, Kong CS, Lipecki J, Fishwick KJ, Odendaal J, Ewington LJ, et al. 2020 Recurrent pregnancy loss is associated with a pro-senescent decidual response during the peri-implantation window. Commun Biol 3 37.

Masuda H, Anwar SS, Buhring HJ, Rao JR & Gargett CE 2012 A novel marker of human endometrial mesenchymal stem-like cells. Cell Transplant 21 2201–2214.

Matsumoto H 2017 Molecular and cellular events during blastocyst implantation in the receptive uterus: clues from mouse models. J Reprod Dev 63 445–454.

Maybin JA & Critchley HO 2015 Menstrual physiology: implications for endometrial pathology and beyond. Hum Reprod Update 21 748–761.

Mills SJ, Cowin AJ & Kaur P 2013 Pericytes, mesenchymal stem cells and the wound healing process. Cells 2 621–634.

Papatheodorou I, Moreno P, Manning J, Fuentes AM, George N, Fexova S, Fonseca NA, Fullgrabe A, Green M, Huang N, et al. 2020 Expression Atlas update: from tissues to single cells. Nucleic Acids Res 48 D77–D83.

Parasar P, Sacha CR, Ng N, McGuirk ER, Chinthala S, Ozcan P, Lindsey J, Salas S, Laufer MR, Missmer SA, et al. 2017 Differentiating mouse embryonic stem cells express markers of human endometrium. Reprod Biol Endocrinol 15 52.

Peyser R, MacDonnell S, Gao Y, Cheng L, Kim Y, Kaplan T, Ruan Q, Wei Y, Ni M, Adler C, et al. 2019 Defining the Activated Fibroblast Population in Lung Fibrosis Using Single-Cell Sequencing. Am J Respir Cell Mol Biol 61 74–85.

Queckborner S, Syk Lundberg E, Gemzell-Danielsson K & Davies LC 2020 Endometrial stromal cells exhibit a distinct phenotypic and immunomodulatory profile. Stem Cell Res Ther 11 15.

Ramachandran P, Matchett KP, Dobie R, Wilson-Kanamori JR & Henderson NC 2020 Single-cell technologies in hepatology: new insights into liver biology and disease pathogenesis. Nat Rev Gastroenterol Hepatol.

Robinson MD, McCarthy DJ & Smyth GK 2010 edgeR: a Bioconductor package for differential expression analysis of digital gene expression data. Bioinformatics 26 139–140.

Saatcioglu HD, Kano M, Horn H, Zhang L, Samore W, Nagykery N, Meinsohn MC, Hyun M, Suliman R, Poulo J, et al. 2019 Single-cell sequencing of neonatal uterus reveals an Misr2+ endometrial progenitor indispensable for fertility. Elife 8.

Satija R, Farrell JA, Gennert D, Schier AF & Regev A 2015 Spatial reconstruction of singlecell gene expression data. Nat Biotechnol 33 495–502.

Schwab KE, Chan RW & Gargett CE 2005 Putative stem cell activity of human endometrial epithelial and stromal cells during the menstrual cycle. Fertil Steril 84 Suppl 2 1124–1130.

Spitzer TL, Rojas A, Zelenko Z, Aghajanova L, Erikson DW, Barragan F, Meyer M, Tamaresis JS, Hamilton AE, Irwin JC, et al. 2012 Perivascular human endometrial mesenchymal stem cells express pathways relevant to self-renewal, lineage specification, and functional phenotype. Biol Reprod 86 58.

Tabula Muris C, Overall c, Logistical c, Organ c, processing, Library p, sequencing, Computational data a, Cell type a, Writing g, et al. 2018 Single-cell transcriptomics of 20 mouse organs creates a Tabula Muris. Nature 562 367–372.

Tempest N, Maclean A & Hapangama DK 2018 Endometrial Stem Cell Markers: Current Concepts and Unresolved Questions. Int J Mol Sci 19.

Yu G, Wang LG, Han Y & He QY 2012 clusterProfiler: an R package for comparing biological themes among gene clusters. OMICS 16 284–287.

