## Supplementary material for "Single cell RNA sequencing redefines the mesenchymal cell landscape of mouse endometrium": Main manuscript

**Supplementary Table 1.** Primary and secondary antibodies with associated working dilutions used to detect proteins in mouse uterine tissues

| **Antibody** | **Supplier and Cat. No. (concentration)** | **Dilution in NGS** |
|---|---|---|
| **Primary antibodies** | | |
| Polyclonal rabbit anti-GFP | Abcam, ab6556 (0.5 ming/ml) | 1/1000 |
| Monoclonal rabbit anti-PDGFRβ | Abcam, ab32570 (0.15 ming/ml) | 1/1000 |
| Monoclonal rabbit anti-CD146 | Abcam, ab75769 (0.123mg/ml) | 1/1000 |
| Polyclonal rabbit anti-NG2 | Abcam, ab12905 | 1/600 |
| Monoclonal mouse anti-αSMA | Sigma, A-2547 | 1/10K |
| Polyclonal rabbit anti-CD31 | Abcam, ab28364 (1mg/ml) | 1/500 |
| Polyclonal rabbit anti-EpCAM | Abcam, ab71916 (1mg/ml) | 1/2000 |
| Polyclonal rabbit anti-MYH11 | Atlas Antibodies HPA015310 | 1/1000 |
| Polyclonal rabbit anti-Desmin | Abcam, ab15200 (1mg/ml) | 1/500 |
| **Secondary antibodies** | | |
| Goat F(ab) anti-rabbit IgG H&L (HRP) | Abcam, ab7171 (1mg/ml) | 1/500 |
| Goat F(ab) anti-mouse IgG H&L (HRP) | Abcam, ab6823 (1mg/ml) | 1/500 |

**Supplementary Table 2.** Flow cytometry antibodies selected and optimised to interrogate mesenchymal cell populations in murine uterus

| Antibody | Supplier Cat. No. (concentration) | Excitation λ (nm) | Emission λ (nm) | Dilution |
|---|---|---|---|---|
| BV421 anti-mouse CD31 | BioLegend 102423 (0.2mg/ml) | 405 | 421 | 1/200 |
| BV421 anti-mouse CD45 | Fischer Scientific BDB560501 (0.2mg/ml) | 404 | 448 | 1/100 |
| APC anti-mouse CD146 | BioLegend 134712 (0.2mg/ml) | 650 | 660 | 1/200 |
| BV605 anti-mouse EpCAM | BioLegend 118227 (0.2mg/ml) | 405 | 603 | 1/400 |
| FITC anti-mouse CD90 | Invitrogen 11-0902-82 (0.5 mg/ml) | 490 | 525 | 1/100 |

**Supplementary Table 3.** Primers and probes used for RTPCR

| Gene name | Accession number | Forward primer sequence | Reverse primer sequence | UPL probe number |
|---|---|---|---|---|
| *Pdgfrβ* | NM_001146168.1 | tcaagctgcaggtcaatgtc | ccattggcagggtgactc | 67 |
| *Mcam* | NM_023061.2 | aaactggtgtgcgtcttcttg | cttttcctctcctggcacac | 27 |
| *Acta2* | NM_007392.2 | ctctcttccagccatctttcat | tataggtggtttcgtggatgc | 58 |
| *Cspg4* | NM_139001 | cttggccttgttggtcagat | cacctccaggtggttctcc | 16 |
| *Myh11* | NM_013607.2 | ttgctgggtttgaaatctttg | ccctctcgctggtactcttc | 1 |
| *Olfr78* | NM_130866 | gcttcctccaaccttctgag | tggagctcgtttcccaaata | 42 |
| *Rgs4* | NM_009062 | tccctcagttaaacaagatgtgc | gtttcatgtcctttgcactcc | 4 |
| *Rgs5* | NM_009063.4 | cagacagaggcccctaaaga | agacggttccaccaggttc | 9 |
| *Vcam1* | NM_011693 | tcttacctgtgcgctgtgac | actggatcttcagggaatgagt | 47 |
| *Kitl* | NM_001347156.1 | cagcgctgcctttccttat | ccttggttttgacaagaggatt | 68 |
| *Kcnj8* | NM_001330363 | gagaaaggcaccatggagaa | ggagaagagaaacgcagacg | 109 |

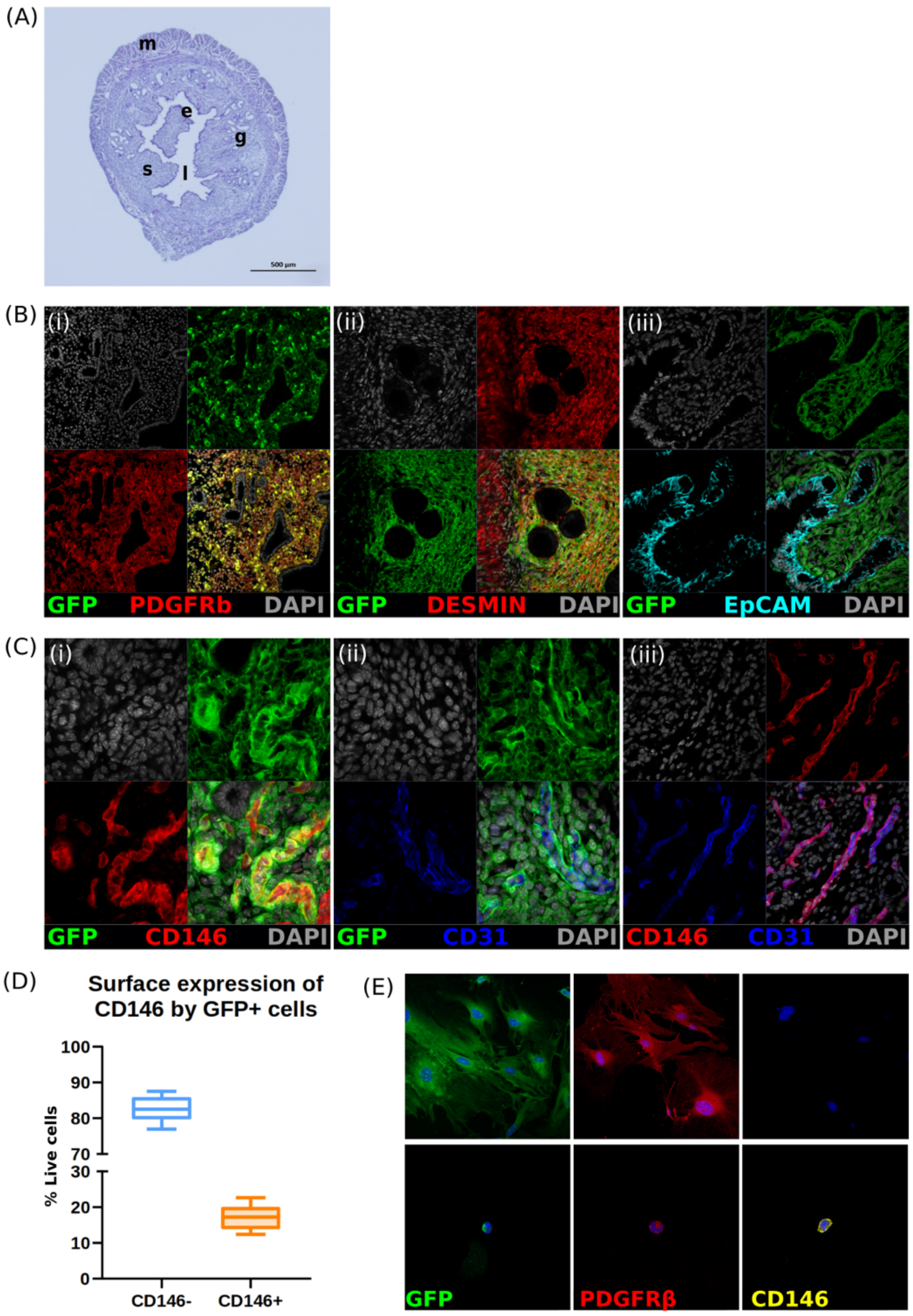


**Supplementary Figure 1. Validation of GFP reporter protein expression by mesenchymal cells in *Pdgfrb*-BAC-eGFP mouse endometrium. (A)** H&E staining: mouse uterus cross section to highlight the outer myometrium (m), endometrial stroma (s), inner lumen (l), luminal epithelium (e) and epithelial glands (g). **(C)** Split channel immunofluorescence showing expression and/or co-expression of: **(i)** GFP and PDGFRβ; **(ii)** GFP and Desmin; **(iii)** GFP and EpCAM in *Pdgfrb*-BAC-eGFP mouse endometrial tissue. **(C)** Split channel immunofluorescence showing expression and/or co-expression of: **(i)** GFP and CD146; **(ii)** GFP and CD31; and **(iii)** CD146 and CD31; in *Pdgfbβ*-BAC-eGFP mouse endometrial tissue. **(D)** Immunofluorescence detection of GFP, PDGFRβ and CD146 on cytospins of $GFP^{dim}$CD146- and $GFP^{bright}$CD146+ cells isolated from *Pdgfrb*-BAC-eGFP mouse endometrium by FACS confirming $GFP^{dim}$ cells to be PDGFRβ+CD146- while $GFP^{bright}$ cells are PDGFRβ+CD146+ (representative images; n=4).

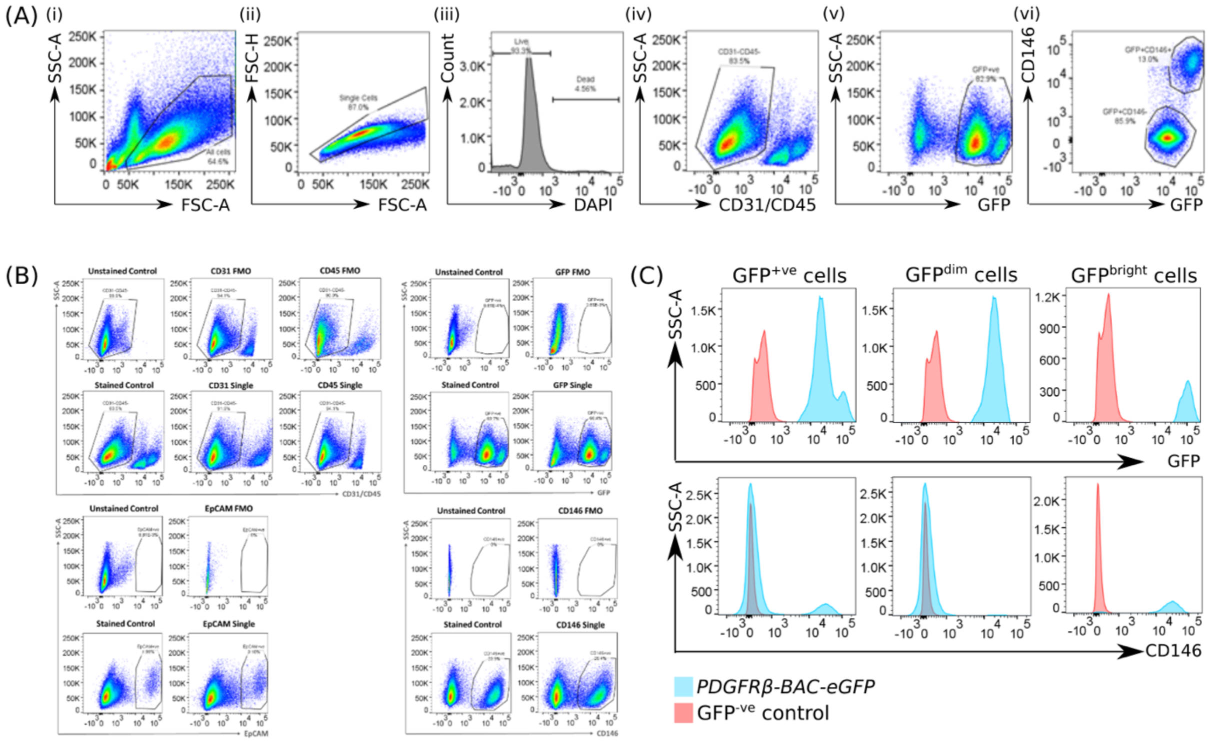


**Supplementary Figure 2. Parameters employed to draw analytical gates to analyse uterine tissues collected from *Pdgfrb*-BAC-eGFP mice by flow cytometry. (A)** Sequential gating strategy employed to exclude and/or analyse GFP/PDGFRβ+ cell populations in mouse uterine tissue: **(i)** All cells; **(ii)** Single cells; **(iii)** Live cells; **(iv)** CD31-CD45- cells; **(v)** GFP+ cells; **(vi)** CD146+/- cells; percentages displayed are a frequency of parent gate. **(B)** Analytical gates to capture cells of interest were all determined using controls respective to each marker: unstained, fully stained, single stains and 'fluorescence minus one' (FMO) controls for CD31, CD45, EpCAM, GFP and CD146. (representative plots, n=4). **(C)** Histograms displaying the expression of GFP and CD146 by GFP+, $GFP^{dim}$ and $GFP^{bright}$ cells in *Pdgfrb*-BAC-eGFP uterine tissues (blue) and C57BL/6 uterine tissue (red) (representative plots; n=8).

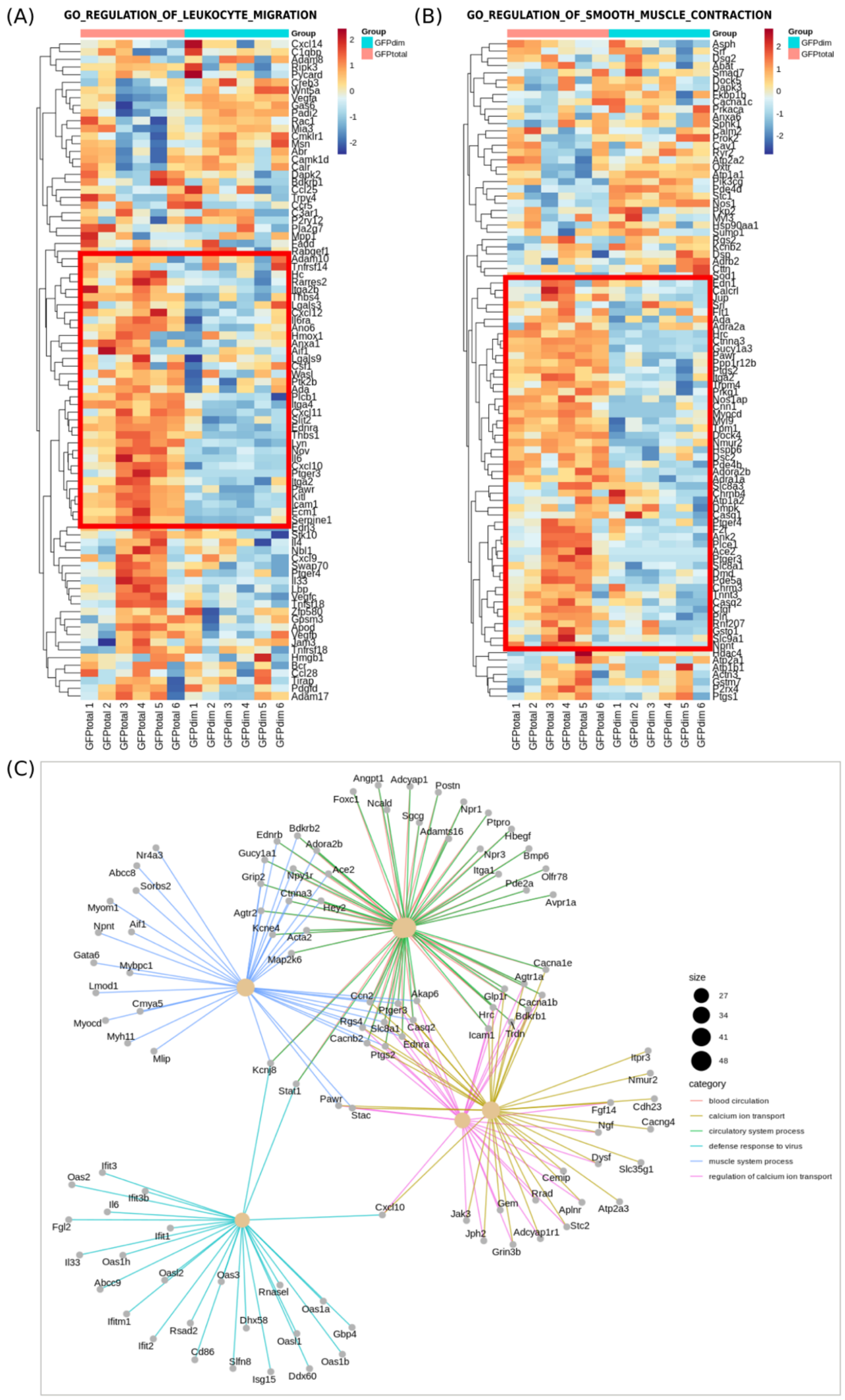


**Supplementary Figure 3. Gene ontology enrichment analysis for biological process (BP) terms enriched by putative pericyte genes identified through bulk mRNA sequencing. (A)** Heatmap: expression of key genes associated with 'regulation of leukocyte migration' across GFP$^{total}$ and GFP$^{dim}$ samples. **(B)** Heatmap: expression of key genes associated with 'regulation of smooth muscle cell contraction' across GFP$^{total}$ and GFP$^{dim}$ samples; red box indicates genes ubiquitously expressed by GFP$^{total}$ samples when compared to GFPdim samples, indicative of expression by GFP$^{bright}$

cells/putative pericytes. **(C)** Gene-concept network (cnet) plot generated using the ‘clusterProfiler’ package to depict the linkages of genes and multiple biological concepts as a network and identify genes contributing to more than one ontology: *Kcnj8*, *Stat1*, *Pawr*, *Stac*, *Cxcl10*, *Ednrb*, *Acta2*; dotsize: number of genes in data associated with each GO term.

(A)

| Gene | Expanded Name | Mean CPM Naive | Mean CPM Stromal | LogFC (>2) |
|---|---|---|---|---|
| *Acta2 (αSMA)* | Actin, alpha 2 smooth muscle | 48.62480465 | 1.059755434 | 5.508426009 |
| *Myh11* | Myosin heavy polypeptide 11 | 112.3382175 | 1.871607608 | 5.915143007 |
| *Cspg4 (NG2)* | Neural glial antigen 2 | 7.623761625 | 0.572417667 | 3.710645725 |
| *Olfr78* | Olfactory receptor 78 | 13.14954306 | 0 | 10.64494967 |
| *Rgs4* | Regulator of G-protein signalling 4 | 30.59502037 | 1.376631718 | 4.463932917 |
| *Rgs5* | Regulator of G-protein signalling 5 | 659.6824379 | 3.541085656 | 7.545825823 |
| *Vcam1* | Vascular cell adhesion molecule 1 | 18.8761171 | 2.328397054 | 3.013912524 |
| *Mcam (CD146)* | Melanoma cell adhesion molecule | 7.13375629 | 0.463077708 | 3.925662267 |
| *Kitl* | Kit ligand | 16.11201615 | 0.682810577 | 4.552554094 |
| *Kcnj8* | Potassium channel, subfamily J, member 8 | 11.43342976 | 0.028477121 | 8.418738739 |

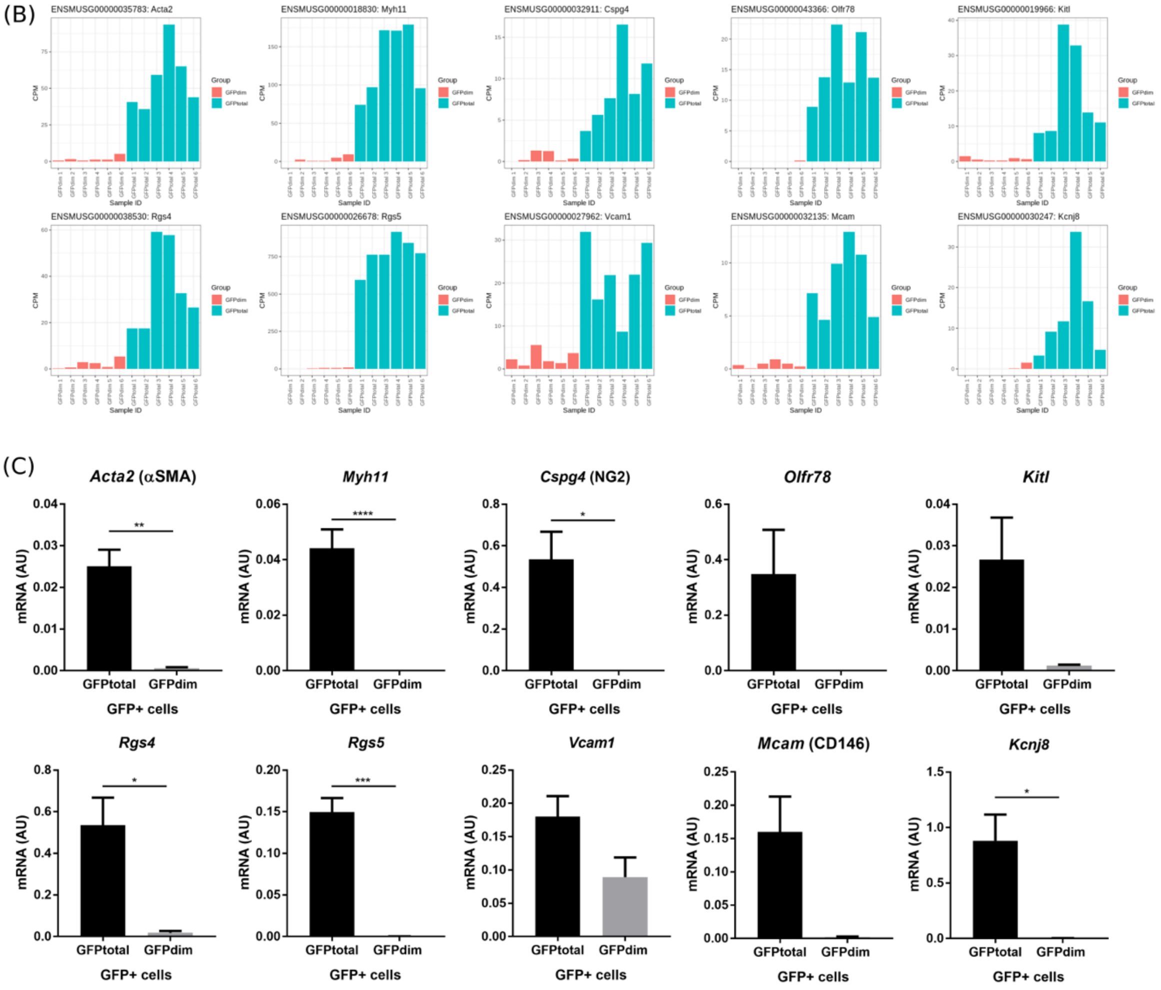


**Supplementary Figure 4. Selection of putative pericyte gene candidates to validate Bulk mRNA sequencing data in mouse uterine tissue samples by qPCR. (A)** Table outlining the mean CPM in GFP$^{total}$ and GFP$^{dim}$ samples and the resultant logFC of DEgenes selected for validation studies. **(C)** Barcharts: expression of selected DEgenes (presented as CPM) across GFP$^{total}$ and GFP$^{dim}$ samples to confirm ubiquitously higher counts in GFP$^{total}$ samples. **(D)** qPCR analysis of the expression of putative pericyte genes in a new set of GFP$^{total}$ cell and GFP$^{dim}$ cell samples isolated from *Pdgfrb*-BAC-eGFP mouse uterine tissue. mRNA for *Acta2*, *Myh11*, *Cspg4* (NG2), *Olfr78*, *Kitl, Rgs4*, *Rgs5*, *Vcam1*, *Mcam* (CD146) and *Kcnj8* was more highly expressed in GFP$^{total}$ cells when compared to GFP$^{dim}$ cell samples (n=4; one-way ANOVA, Holm-Sidak's multiple comparisons test,*p<0.05, **p<0.01, ***p<0.001, ****p<0.0001).

(A)

| Summary Statistics (cellranger_count) | |
|---|---|
| **Cells** | |
| Estimated number of cells | 6,379 |
| Fraction reads in cells | 94.3% |
| Mean reads per cell | 71,067 |
| Median genes per cell | 1,847 |
| Total genes detected | 19,268 |
| Median UMI counts per cell | 4,316 |
| **Sequencing** | |
| Number of reads | 453,334,959 |
| Valid barcodes | 97.7% |
| Valid UMIs | 100.0% |
| Sequencing saturation | 85.1% |
| Q30 bases in barcode | 94.4% |
| Q30 bases in RNA read | 87.0% |
| Q30 bases in sample index | 92.6% |
| Q30 bases in UMI | 94.1% |
| **Mapping** | |
| Reads mapped to genome | 89.7% |
| Reads mapped confidently to genome | 85.6% |
| Reads mapped confidently to intergenic regions | 4.2% |
| Reads mapped confidently to intronic regions | 24.4% |
| Reads mapped confidently to exonic regions | 56.8% |
| Reads mapped confidently to transcriptome | 54.4% |
| Reads mapped antisense to gene | 1.1% |

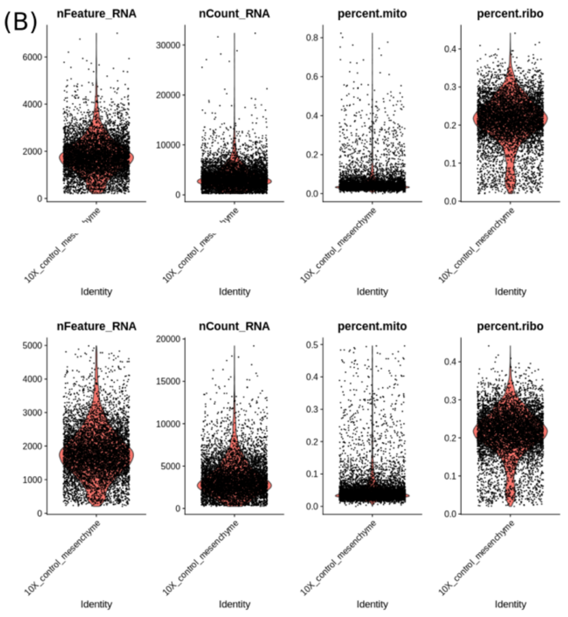


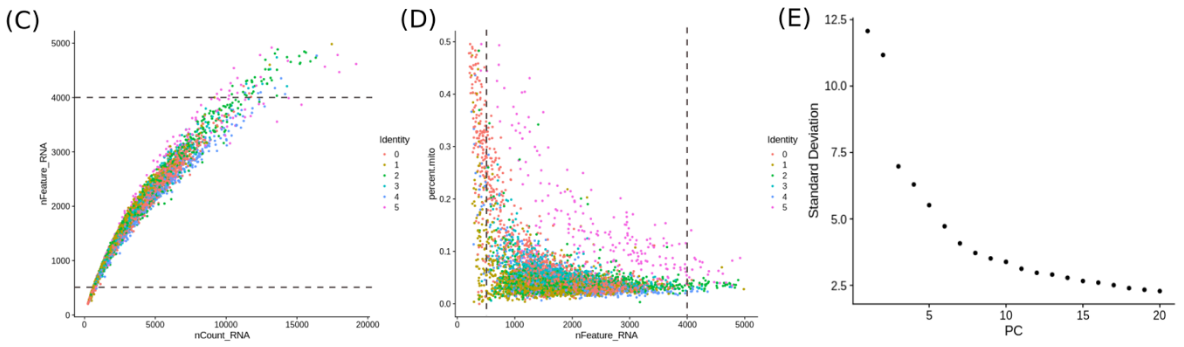


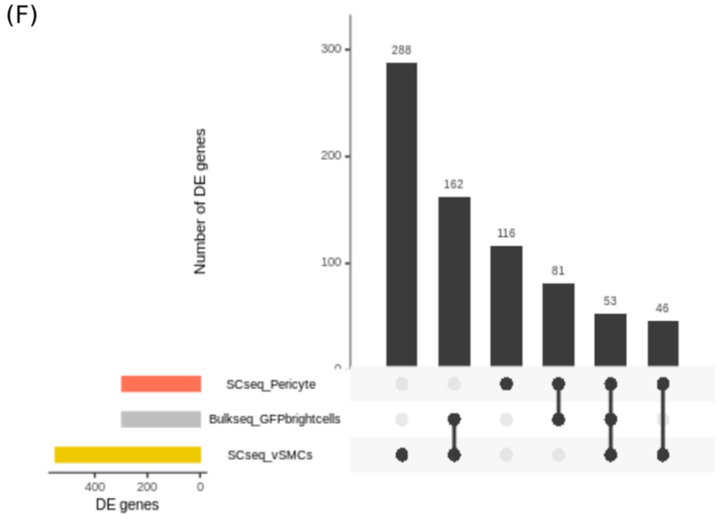


**Supplementary Figure 5. Quality control of the single-cell RNA sequencing reads prior to downstream analyses. (A)** Summary statistics of the counts matrix resulting from the cellranger_count analysis pipeline. **(B)** Cell QC based on three QC covariates: violin plots displaying number of counts per barcode (count depth; nFeature_RNA), number of genes per barcode (nCount_RNA), the fraction of counts from mitochondrial genes per barcode (percent.mito) and the fraction of counts from ribosomal genes per barcode (percent.ribo) before filtering (top panel) and after filtering outlier peaks by thresholding: nFeature_RNA >200 & < 4000, percent.mito <0.5 and percent.ribo <0.5 (bottom panel).

**(C)** Number of genes (nCount_RNA) versus number of counts (nFeature_RNA): cells where detection of number of genes is high have high counts indicative of doublets. These cells are filterd out by applying thresholds displayed by dashed lines. **(D)** Number of genes (nFeature_RNA) versus the percentage of mitochondrial genes (percent.mito) coloured by cell cluster: mitochondrial read fractions are only high in particularly low count cells with few detected genes. These cells are filtered out by applying thresholds displayed by dashed lines. **(E)** Scree plot: plot of the eigenvalues of the first 20 principal components of the data used to determine the number of principlal components to keep in a PCA: the 'elbow' of the graph reveals that the first 10 principal components should be retained as significant and used in downstream analyses. **(F)** Upset plot: visualisation of the intersection between the gene list generated by bulk mRNA seq of $GFP^{bright}$ cells (grey bar) and those generated by scRNAseq for perivascular cell subpopulations (vSMCs: yellow bar; pericytes: orange bar).
